# SMC protein RecN drives RecA filament translocation and remodelling for *in vivo* homology search

**DOI:** 10.1101/2021.08.16.456443

**Authors:** Afroze Chimthanawala, Jyotsana J Parmar, Sujan Kumar, Krishnan S. Iyer, Madan Rao, Anjana Badrinarayanan

## Abstract

While the molecular repertoire of the homologous recombination pathway is well-studied, the search mechanism that enables recombination between distant homologous regions is poorly understood. Earlier work suggests that the recombinase RecA, an essential component for homology search, forms an elongated filament, nucleating at the break site. How this RecA structure carries out long distance search remains unclear. Here, we follow the dynamics of RecA after induction of a single double-strand break on the Caulobacter chromosome. We find that the RecA-nucleoprotein filament, once formed, rapidly translocates in a directional manner in the cell, undergoing several pole-to-pole traversals, until homology search is complete. Concomitant with translocation, we observe dynamic remodelling of the filament. Importantly in vivo, the RecA filament alone is incapable of such long distance movement; both translocation and dynamic remodelling are contingent on action of SMC-like protein RecN, via its ATPase cycle. We provide a stochastic description of RecN-regulated changes in filament length during translocation via modulation of RecA assembly-disassembly. In summary, we have uncovered the three key elements of homology search driven by RecN: mobility of a finite segment of RecA, filament remodelling and ability to conduct multiple pole-to-pole traversals, which together point to a novel optimal search strategy.

## Introduction

DNA double-strand breaks (DSBs), if left unrepaired (or incorrectly repaired) can lead to loss of genetic information, chromosomal rearrangements, mutagenesis and even cell death. Homologous recombination, a process that usually ensures error-free repair of DSBs has emerged as an evolutionarily conserved pathway from bacteria to eukaryotic cells^1,2^. Recent work has shown that recombination-based repair is not only local, such as during post-replicative cohesion, but that even distant homologous regions can participate in such repair^2–12^. This is particularly relevant for bacteria, where an extensive post-replicative cohesion period is absent^13–19^. Indeed, cells can face DSBs even outside the context of replication (examples include X-rays, IR radiation, chemicals and metabolites such as bleomycin, Topoisomerase dysfunction (from inhibitor action for example), processing of closely-spaced CPDs, endonucleases produced by the cell and bacterial toxins that directly cause DSBs^7,8,20–25^). Several of these breaks rely on recombination-based repair^7,8,21,26,27^. Given the potential lethality associated with unrepaired breaks^7,9,20,21,27–32^, it is vital that there be regulatory mechanisms to sense and initiate, translocate and search and finally identify even distant homology regions for repair. These strategies for long distance homology search and repair must potentially depend on the spatial organisation of chromosomes - its extent, geometry and dimensionality^33^ - and must involve energy transduction.

In bacteria, for instance, an extensive homology search programme is effected by the highly conserved recombinase, RecA^34–36^. In the first step of the recombination pathway, DSB-ends are processed to reveal ssDNA overhangs, on which RecA can assemble into a filamentous structure^4,37,38^ which in turn initiates homology search. *In vitro* studies have shown that RecA molecules in the nucleoprotein filament possess ATPase activity that can influence (a) formation of a continuous filament during RecA loading on ssDNA^39^, (b) growth and shrinkage of the filament^40–42^, (c) release of the filament from heterologous pairing^35,43^, and (d) RecA turnover during strand invasion^42–44^. *In vivo* studies on double-strand break repair have made use of engineered endonuclease-based DSB-inducing systems (such as the I-SceI system) to make targeted breaks on the chromosome and track the associated dynamics^4–6,11,12,45–47^. Using such a system in *E. coli* to study dynamics of long-distance homology search, imaging of RecA has revealed the presence of large assemblies of RecA, described as RecA *filaments* (or ’bundles’), that can even span the length of the cell^5,6^. How this RecA structure carries out homology search is unknown.

In addition, several accessory proteins are reported to modulate the association and turnover of RecA with ssDNA^48,49^. Of these, RecN, a highly conserved repair-associated protein, has been shown to interact with RecA and is essential for DSB repair^49–52^. RecN belongs to the Structural Maintenance of Chromosome (SMC) family of proteins, that play central roles in chromosome dynamics^53^. While their function in chromosome organization and segregation in bacteria has been well-characterized, their role in DNA repair pathways has not been clearly established. Previous studies have suggested a role for RecN in homologous recombination, either via its influence on kinetics of strand-invasion by the RecA filament or by facilitating global chromosomal cohesion between replicated sisters^49,51,54,55^. Indeed the potential relevance of RecN in recombination-based repair is underscored by its extensive conservation with other core recombination-associated proteins such as RuvABC and RecA across bacterial genomes, as well as across domains of life^56^.

In conjunction with other molecular regulators, homology search appears to be an efficient and robust process; recent studies reveal that long-distance search and repair can typically be completed within a single generation time of bacterial growth^3,4,6^. How RecA orchestrates this search, the dynamics of the nucleoprotein filament during this process, and the role for RecA regulators in facilitating or modulating homology search, remain largely unknown. To elucidate the mechanism of homology search *in vivo,* we use quantitative live cell imaging to track RecA during DSB repair in the bacterium *Caulobacter crescentus,* together with a theoretical description of RecA dynamics during search.

Here we present evidence for a novel search mechanism involving a RecA nucleoprotein filament that translocates in a directional manner from the break site located at one cell pole, to the opposite cell pole. Unexpectedly, the RecA filament simultaneously undergoes dynamic changes to its structure, while traversing back-and-forth across the length of the cell before finding its homologous partner and initiating homology pairing. Both, large scale movement and dynamic remodelling of RecA are contingent on its association with RecN. We present a stochastic description, where RecN influences dynamic changes in RecA filament lengths via regulation of rates of RecA subunit assembly-disassembly. Such dynamical remodelling and directional movement of the RecA nucleoprotein filament points to a robust mechanism of iterative search, thus enabling recombination between distant sites of homology.

## Results

### *In vivo* imaging of RecA reveals key steps of homologous recombination

To follow RecA-dependent search dynamics *in vivo,* we use the I-SceI system developed in *Caulobacter crescentus*^4,38^ that allows us to visualize DSB repair between distant homologous regions (Fig. S1A). Briefly, we insert a unique restriction site, *I-SceI,* near the replication origin, which is usually polarly localized in the cell. We induce a DSB by regulated expression of the endonuclease, I-SceI, that cleaves the site to generate a break. We further marked a region of the chromosome near the break-site using fluorescently-tagged MipZ that associates with the origin region. DSB induction results in loss of localization of the MipZ marker close to the break (due to processing of DNA by the helicase-nuclease complex AddAB^4,38^) serving as a reliable proxy to identify cells where only a single copy of the chromosome is cleaved (with only one MipZ marker now being visible). Restoration of the second marker is indicative of repair. Such complete repair events, as reported previously^4^, are observed in 30% of all imaged cells during the course of a single experiment (with 22% cells losing both MipZ foci (suggesting two breaks), 18% cells facing no breaks and 30% cells not having completed repair within the imaging window). Using this extensively characterized system^4,38^, we follow the process of homology search in two distinct cell types, both blocked for new rounds of replication initiation: pre-divisional cells with two completely replicated and segregated chromosomes (to follow homology search and subsequent recombination-mediated repair of distant homologous regions) and non-replicating swarmer cells with a single chromosome (to follow homology search, without the presence of the homologous repair template)^4,38^.

In both cell types, we monitor the dynamics of RecA using live cell imaging of RecA-YFP. Given that endogenously-tagged RecA perturbs the functioning of the protein, we use a well-described strategy in *E. coli,* where RecA is expressed ectopically in a wild type RecA background^4–6,51,57,58^. This strain, as previously described in *Caulobacter*^4^, results in RecA-YFP levels being 30-50% of wildtype RecA levels and is found to be comparable to the wild type background in terms of function in generic DNA damage repair and recombination-mediated DSB repair (as assessed via CFU survival measurement on generic as well as DSB-inducing damage). Furthermore: a. RecA-YFP or RecA-mCherry do not localize to the break site in the absence of damage (Fig. S4F) and localize to DSBs only after break processing by AddAB ^4^. This is in contrast to RecA tags in *E. coli* that can localize polarly even in the absence of damage ^5,51,57^ b. RecA localization persists only until damage repair. Upon DSB repair (re-emergence of 2 MipZ foci), RecA does not localize in a discrete manner in the cell. This suggests that RecA localization is associated with damage. (Fig. 1A, n=100cells). c. RecA localizes at the site-specific break only. We find that RecA localization correlates with the spatial location of the break site within the cell. (Fig. S2 representative from n=100 cells in each example). d. RecA and RecN colocalize. Both proteins localize only in the presence of DNA damage (Fig. S3A-B, S4F) and we observe them to be colocalized (Fig. S3C). e. Read-out of SOS response (using MMC or Norfloxacin treatment, following previously-described protocols in *Caulobacter*) shows that cells carrying the fluorescently-tagged RecA construct induce the response similar to wild type conditions (Fig. S1D).

**Fig. 1.**
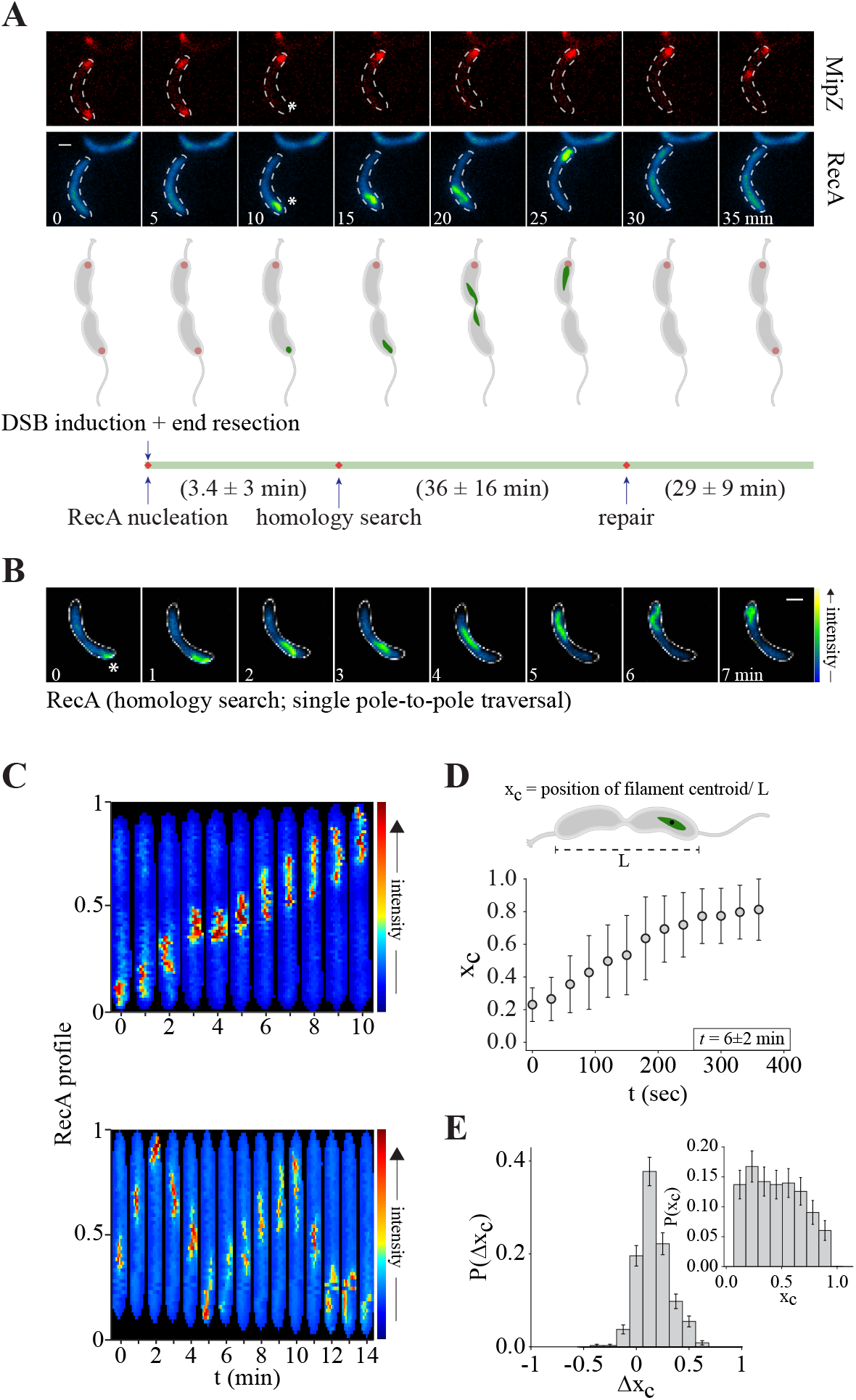
Dynamics of RecA filament during homologous recombination. (A) Montage of key events of homologous recombination observed via live-cell imaging of both break/ homology site (MipZ; red) as well as DSB dynamics (RecA; green). Top panel shows representative cell examples for each stage (t0 refers to the first time point of imaging; bottom panel provides quantitative estimation of duration of the key events described across multiple cells (n = 100 repair events). Single DSB induction (white asterisk) is marked by the disappearance of one of the two MipZ localizations. *RecA nucleation* is defined as the period between DSB induction and the initiation of RecA localization at the break site, followed by *homology search,* marked by the translocation of the RecA filament in the cell. *Repair* is marked by the loss of RecA filament after search, followed by the reappearance of two MipZ markers. The time line below shows the period of RecA nucleation and loading (after break induction), homology search and repair (n = 100). Scale bar: 2 μm here and in all other images. (B) Montage of RecA-YFP tracked over time (duration of a single pole-to-pole traversal event) after induction of a single DSB during homology search (white asterisk). Here, *t0* is defined as the time point prior to the observation of a positive displacement of the RecA localization from the pole at which it first localized (DSB pole) (C) Representative RecA-YFP fluorescence profiles across [top] one pole-to-pole traversal and [bottom] multiple traversals. (D) Centroid position of the RecA filament x_*c*_ (measured relative to cell length) versus time over a single traversal. Plot shows mean and standard deviation from data collected over n = 99 cells. Mean traversal time *t* = 6 ± 2 min is indicated. (E) Bar graph representing the probability distribution of the centroid steps Δx_*c*_, (i.e. the algebraic difference between the centroid positions at two consecutive time points (60 sec apart)), shows a bias towards positive values and a weight at Δx_*c*_ = 0 (Error bars represent sample error). [inset] Bar graph representing the probability distribution of x_*c*_ taken over a single traversal cycle (n = 795). All data shown are from pre-divisional cells.

We find that *Caulobacter* RecA-YFP forms discrete structures (foci or elongated filaments) only in the presence of DNA damage. In the present study, we use the term ‘filament’ to describe diffraction-limited RecA localizations. Whether this includes bundles or single filament of RecA polymers^5,6^ remains to be determined. In the absence of damage, fluorescence is diffuse throughout the cell (Fig. S4F). We observe three distinct stages following a DSB near the cell pole – (i) *nucleation and growth* of RecA filament near the pole where MipZ localization, proximal to the origin^4,38^, is lost (indicating occurrence of a DSB), (ii) *homology search*, where the RecA filament remodels and moves directionally across the length of the cell (expanded in the next sections), and (iii) *repair,* associated with RecA dissociation followed by the reappearance of two distinct MipZ markers (Fig. 1A; top panel shows representative MipZ and RecA images corresponding to each stage (in this case t0 refers to the first time point of imaging. Cells are imaging until ‘repair’ is complete and two distinct MipZ markers are now visible) and the bottom panel provides quantitation of duration of each stage described, from multiple cells estimated from n = 100 complete repair events).

### Growth and directional movement of RecA filament during homology search

We now characterize the dynamics of RecA during the homology search phase. In contrast to the static structures previously reported in *E. coli*^5,6^, we find that once RecA nucleates at the DSB site near the pole, it rapidly grows into an elongated filament, following which it dynamically moves across the length of the cell (Fig. 1B, C, S2A, S2D; *Supplementary Video 1*) We follow the time series of the centroid position x_*c*_(t) of the RecA filament along the long-axis of the cell, relative to the cell length L (while analysing traversals, *t0* is defined as the time point prior to the observation of a positive displacement of the RecA localization from the pole at which it first localized (DSB pole); see *Supplementary Results* for details of quantitative analysis). From its initial position x_*c*_(0) close to the nucleation site, we find that the centroid progresses in a stochastic but biased manner, towards the opposite cell pole (Fig. 1D,E). Having reached the opposite cell pole, the RecA filament starts translocating in the reverse direction (Fig. 1C, S2A). Defining a pole-to-pole translocation as a *traversal*, we find that the mean traversal time of the RecA filament is 6 ± 2 min, and typically undergoes multiple traversals (mean = 4) before homology pairing (Fig. 1D).

On average, the centroid position is uniformly distributed across the cell length (Fig. 1E, inset). This indicates that there are no hot-spots within the cell that specifically regulate RecA filament dynamics. The distribution of the centroid steps Δx_*c*_, the algebraic difference between the centroid positions at two consecutive time points, exhibits a bias towards positive values, indicating a systematic movement of the RecA filament in one direction. The nonzero weight of this distribution at Δx_*c*_ = 0 is an indication that the RecA filament often stalls (Fig. 1E).

Significantly, we observe that when the DNA break is made midcell (+780kb from the origin)^4^, the RecA filament nucleates near midcell, and moves towards *either one of the cell poles* (Fig. S2B). Similarly, in non-replicating swarmer cells with a single copy of the chromosome, the RecA filament seems to exhibit interminable pole-to-pole traversals (Fig.2F, S2C; Supplementary Video 2). These observations imply that the homology search dynamics proceeds independent of the homologous repair template. Interestingly, although there are two break ends upon DSB induction, we rarely observed two distinct RecA filaments. Thus, RecA appears to form a single filamentous structure spanning both break ends, which translocates across the cell during homology search.

**Fig. 2.**
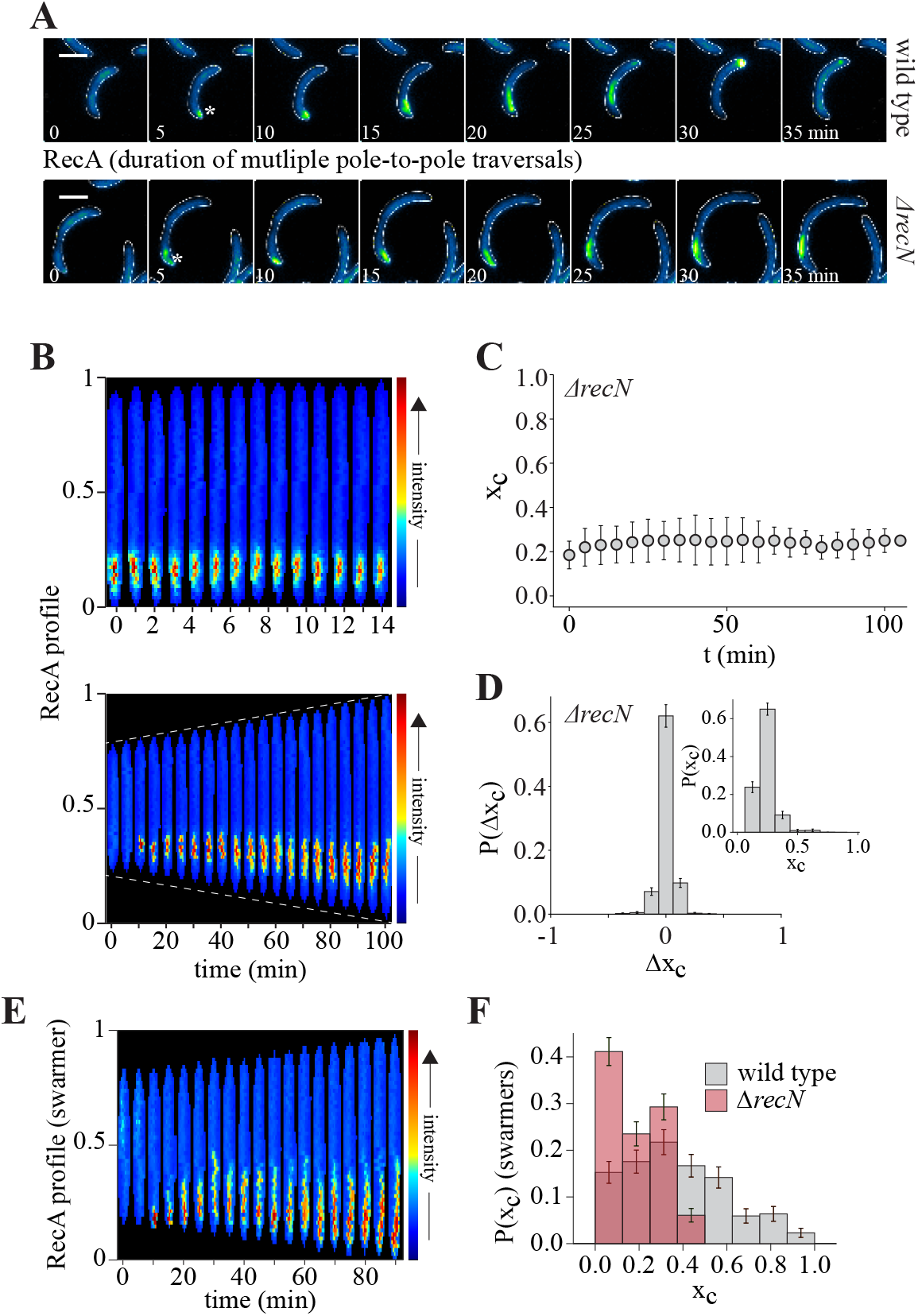
RecN is essential for RecA filament movement *in vivo*. (A) Montage of RecA-YFP tracked over multiple pole-to-pole traversal events after induction of a single DSB (white asterisk) - [top] in wild type, [bottom] in cells lacking *recN* (for the same duration as wild type cells above). Here, *t0* is defined as the first time point when a RecA localization is observed. Examples shown from pre-divisional cells. (B) Representative RecA-YFP fluorescence profiles in a *recN* deletion background. [top] RecA is tracked for the expected duration of two pole-to-pole traversals; [bottom] over expected duration of multiple traversals. Data shown are from pre-divisional cells. (C) Centroid position of the RecA filament x_*c*_ versus time for cells lacking *recN.* Plot shows mean and standard deviation from data collected over n = 50 cells. Data shown are from pre-divisional cells. (D) Probability distribution of the centroid steps Δx_*c*_ shows no bias (Error bars represent sample error). [inset] Probability distribution of x_*c*_ taken over all time points (n = 852), in cells lacking *recN.* Data shown are from pre-divisional cells. (E) Representative RecA-YFP fluorescence profile in a *recN* deletion background. DSB is induced in non-replicating swarmer cells with a single chromosome, thus lacking a homologous template for repair. (F) Probability distribution of x_*c*_ in wild type (n = 911) and cells lacking *recN* (n = 1039). DSB is induced in non-replicating swarmer cells with a single chromosome, thus lacking a homologous template for repair.

### RecN is essential for movement of the RecA filament *in vivo*

We now inquire into the molecular basis of the dynamics of the RecA filament during homology search and for holding break ends together during filament translocation. An essential and highly conserved member of the bacterial recombination pathway is the SMC-like protein, RecN^50,52^. However, it still remains unclear as how RecN influences RecA-mediated homologous recombination *in vivo*. Given the universal role of SMC proteins in organizing DNA (for example, via DNA bridging or DNA loop extrusion^53^, we asked whether RecN contributed to RecA dynamics and/ or organization of break ends specifically during homology search.

Consistent with reports in other bacterial systems^50,52^, we find that RecN is essential for successful DSB repair in *Caulobacter.* In absence of *recN* we record no detectable repair events, together with an increased sensitivity to DSB-inducing agents (Fig. S3E–F, S4G). We use the bacterial-two-hybrid system to show that *Caulobacter* RecA and RecN physically interact with each other (Fig. S3D). We also find that the recruitment of RecN to damage sites is RecA dependent (and not only on SOS-dependent expression) (Fig. S3A, B). Furthermore, RecA and RecN are colocalized at a DSB site, with no detectable instances of RecN localizing away from the RecA structure. However, we do observe examples of RecN localizing towards one end of an elongated RecA filament (Fig. S3C).

Given these observations, we investigate how RecN impacts RecA dynamics and homology search. For this, we first delete *recN* and follow the dynamics of RecA during the search process via live-cell imaging. As in case of wild type, we still observe a single RecA localization in *recN* cells, suggesting that RecN does not contribute to holding break ends together. Instead, we find that upon *recN* deletion, the RecA filament ceases to translocate in pre-divisional cells (Fig. 2A-D; *Supplementary Video 3*) (here, *t0* is defined as the first time point when a RecA localization is observed). Unlike wild type, the RecA filament remains rooted to the pole where the break occurred (Fig. 2B, C, D (inset)) and does not undergo pole-to-pole traversal over the entire course of imaging, up to 100 min (Fig. 2C). Concomitantly, no RecA dissociation is observed, consistent with the idea that recombination between distant homologous regions involves RecN-driven RecA movement. In our experiments, all repair events are associated with the formation and mobility of RecA filaments (and search across the cell). Lack of search (as seen with the absence of RecN) results in 0 repair events (Fig. S4G).

We find a similar effect on the RecA filament translocation in the absence of *recN*, in non-replicating swarmer cells with only a single copy of the chromosome, in support of our assertion that RecN-driven RecA movement for homology search occurs independent of a homologous template for repair (Fig. 2E, F; *Supplementary Video 4*). Finally, these perturbations to RecA dynamics are observed only in cells lacking the repair-specific SMC-like protein (RecN) and not the SMC protein (Smc) involved in chromosome organization and segregation in non-damage conditions^53^ (Fig. S3F, G).

To study which aspects of RecA activity is perturbed in the absence of *recN,* we assess the effects of *recN* deletion on kinetic rates at different stages of the RecA dynamics. In the nucleation stage, we measure the time taken from MipZ disappearance to the appearance of a RecA nucleation event at a DSB site, and find it comparable to wild type cells (Fig. S4A). In the growth stage following nucleation (and prior to search), we find that the initial rates of RecA loading are unaltered in *recN* deleted cells (Fig. S4D, E). In line with this, we observe that the frequency of RecA nucleation events across *recN* cells is comparable to wild type, even under generic mitomycin-C (MMC) damage (Fig. S4F).

Finally, we examine whether the absence of RecA filament movement in *recN* deleted cells is a consequence of a RecA filament instability. Previous studies have shown that cells lacking *recA* or with mutants of RecA that are unable to stably associate with a DSB, undergo ‘rec-less’ degradation, resulting in excessive loss of DNA around a break site^34,59^. Hence we measure DNA degradation in cells with a single DSB (+780kb from the origin in non-replicating swarmer cells) in wild type, *recA* and *recN* backgrounds, via a deep-sequencing assay^38,60^. Our results show that cells lacking *recN* have degradation profiles that overlap with that of wild type (Fig. S4B, C), while cells lacking *recA* show extensive loss of DNA around the break site. In addition, *recN* deleted cells do not have perturbed SOS induction (Fig. S4H). These observations suggest that RecA filament association with a DSB is not affected in the absence of *recN,* although its movement during homology search is stalled.

### RecN-mediated RecA translocation is accompanied by dynamic remodelling of the filament during homology search

Together with the observation that RecN does not affect RecA filament formation at a break site, we find that the average filament lengths are comparable in both wild type and *recN* cells. The length of the RecA filament is typically a small fraction of the cell length and does not extend over the entire cell. In wild type, filament mean fractional length is 〈*l*〉 = 0.2 ± 0.1 (Fig. 3E), that corresponds to 1.0 ± 0.5μm for cell lengths that range between 5 - 6μm. In case of *recN* deleted cells, mean fractional length is 〈*l*〉 = 0.2 ± 0.06, that corresponds to 1.2 ± 0.5μm for cell lengths that range between 7 - 9μm (*Supplementary Results*). However, although the mean filament lengths are similar, we note that the distribution of Δ*l*, the algebraic difference between the filament lengths at two consecutive time points shows a large range in case of wild type cells (Fig. 3A). This suggests that the filament is dynamically remodelling during translocation, both of which appear to be compromised in the absence of *recN* (Fig. 2D, 3A).

**Fig. 3.**
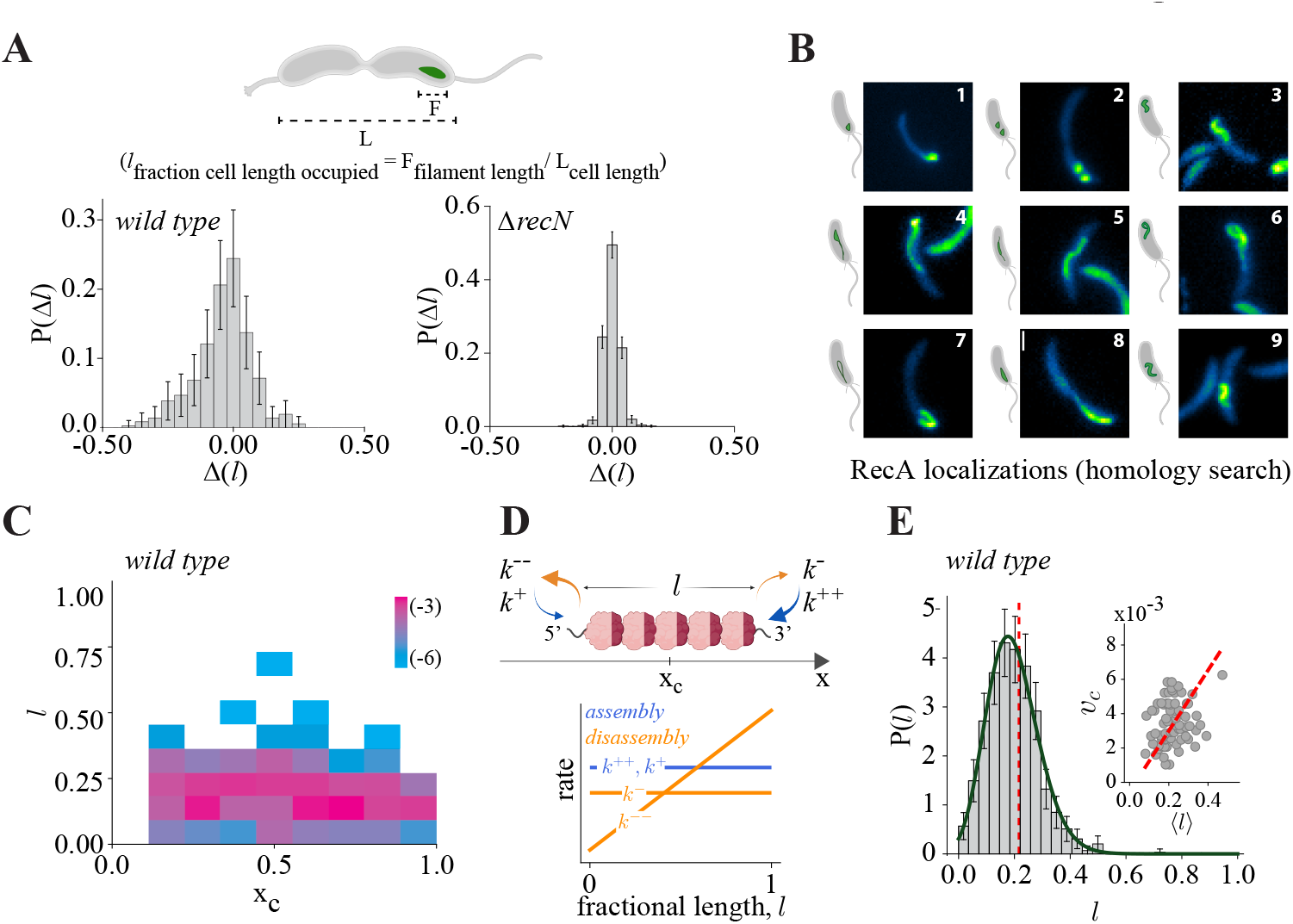
RecN-mediated RecA translocation is accompanied by dynamic remodelling of the filament during homology search. (A) Probability distribution of the length changes Δ*l*, (i.e. the algebraic difference between the filament lengths at two consecutive time points (60 sec apart)) in wild type (n = 347) and cells lacking *recN* (n = 750). Error bars represent sample error. Data shown are from pre-divisional cells. (B) Representative images of the types of RecA localizations observed during a wild type traversal, with (1, 2) showing *puncta* and (3 - 9) showing *extended filamentous* structures. Data shown are from pre-divisional cells. (C) Joint probability distribution of *l* and x_*c*_ taken over a single traversal in wild type cells (n = 795). Data shown are from pre-divisional cells. (D) (top) Stochastic kinetic description of RecN-assisted dynamics of RecA filament in terms of its centroid position and length measured in the lab frame. The rates of assembly and disassembly at the two ends of the filament are shown. (bottom) To maintain a finite mean length, the rates of disassembly must exceed the assembly rates at large *l* (*Supplementary Results*). Choosing the disassembly rate *k”*to be linear in *l*, and the other rates being constant, allows us to match analytically derived length distribution to the experimentally measured one. (E) The analytically derived distribution P(*l*) (green) (*Supplementary Results*) is compared with the experimental data for wild type (plotted as a density histogram) (fit parameters presented in the *Supplementary Results*). [inset] The theory predicts a relation between the centroid velocity *v_c_* and mean filament length 〈*l*〉. Dashed red line highlights the relationship between *v_c_* and 〈*l*〉, estimated Pearson correlation coefficient r = 0.236 (p-value = 0.045, 95% CI [0.00597689, 0.441833]) (for this estimate we have excluded outliers).

Indeed, in wild type cells, we find that directional movement of RecA during homology search is accompanied by dynamic changes in the RecA filament over time. We observe various types of RecA structures in the cell during a traversal, ranging from a focus to extended filamentous structures (Fig. 3B). Importantly, we find no correlation between the length of the RecA filament and its centroid position in the cell (Fig. 3C). Thus, RecN-assisted movement of the RecA filament is accompanied by a stochastic remodelling of its length.

How does RecN regulate RecA filament dynamics? The stochastic remodelling of RecA filament length during homology search suggests to us that RecN action must influence assembly-disassembly rates of RecA molecules on the DNA strand (akin to treadmilling) during translocation. Thus, to understand the contribution of assembly-disassembly rates on the observed RecA filament lengths during translocation, we resort to a quantitative stochastic description of the mechanochemistry of RecN-assisted RecA filament dynamics (*Supplementary Results*).

In general, the stochastic kinetic model is parameterized by four kinetic rates acting at the ends of the RecA filament (Fig. 3D): a binding rate *k^+^* at the left end, a binding rate *k^++^* at the right, an unbinding rate *k*^-^ at the right, and a RecN-dependent unbinding rate *k^--^* at the left (noting that the polarity of the filament (and RecA ATPase states) that dictates directionality and assembly-disassembly of RecA molecules^35,40,48,61–63^).

We first note that *in vivo,* the levels of RecA (as previously reported^5^ and Fig. S4E) and ATP^64^ are likely not limiting. With this in mind, if the assembly-disassembly rates are taken to have fixed values, then they cannot be made to balance each other robustly at a particular length and will not give rise to a finite mean filament length at steady state as is observed for wild type (Fig. 3E). One needs a *negative feedback* between the assembly-disassembly rates and the filament length (Fig. 3D, *Supplementary Results*) in order to provide a homeostatic balance leading to a finite mean filament length with small fluctuations about it^65^. We solve this stochastic kinetic model with length dependent rates analytically using a Langevin dynamics with multiplicative noise (*Supplementary Results*). We see that the net bias of the filament to move to the right (Fig. 1E), is contingent on two key features (*Supplementary Results*) - (i) the kinetics rates must violate detailed balance and must thus be driven by energy consumption and (ii) there is a net breaking of left-right symmetry in the binding-unbinding rates, i.e. the net binding at the right end of the filament is larger than the net unbinding at the left end (in the pure assembly-disassembly limit, this is akin to treadmilling^66^).

We obtain closed form expressions for the steady state distribution of the filament length P(*l*) (*Supplementary Results*), which we compare with the observed distribution for the wildtype cells (Fig. 3E). With the disassembly rate *k*^--^ varying with length as in Fig. 3D, we find that the analytically derived steady state P(*l*) is a perfect match to the observed distribution (Fig. 3E). In addition, we predict a linear correlation between the mean centroid velocity and mean filament length at steady state, which appears consistent with the wild type data (Fig. 3E (inset)). Thus, our stochastic theory provides a framework for how RecN-mediated regulation of assembly-disassembly rates of RecA subunits within a filament can result in the filament length dynamics observed during homology search.

### RecN ATPase cycle regulates RecA filament remodelling

While the molecular mechanism of filament remodelling and its impact on translocation warrants future investigations (see below), the following additional observation underscores a central role for RecN in modulating RecA filament lengths: A fundamental characteristic of SMC family proteins is their ATPase cycle, that enables them to influence the organization of the substrates they are bound to as well as translocate across large distances within the cell^67–69^. We hence asked how the RecN ATPase cycle regulates RecA filament dynamics, thus facilitating recombination between distant homologous regions. For this, we generate two mutants of RecN in the conserved WalkerA and WalkerB motif that are likely to result in specific perturbation to either ATP binding or ATP hydrolysis properties of RecN^67,70^: *recN^D472A^* (ATP-binding impaired) and *recN^K35A/E473Q^* (ATP-hydrolysis impaired)^71^. We find that both mutants are compromised in DNA damage repair, similar to the *recN* deletion (Fig. 4C). We next follow the dynamics of RecA during homology search after induction of a DSB in cells carrying either the *recN^D472A^* or *recN^K35A/E473Q^* mutations. We find that these mutants also result in static RecA filaments that are unable to translocate across the cell for homology search (Fig. 4A, B, D-F; *Supplementary Video* 5-6). While translocation of RecA is similarly impaired in both mutants, we notice that RecA filament lengths are differently affected. In case of *recN^D472A^,* RecA filament lengths are comparable to wild type, while they are significantly shorter in the putative RecN ATP hydrolysis mutant (Fig. 4G).

**Fig. 4.**
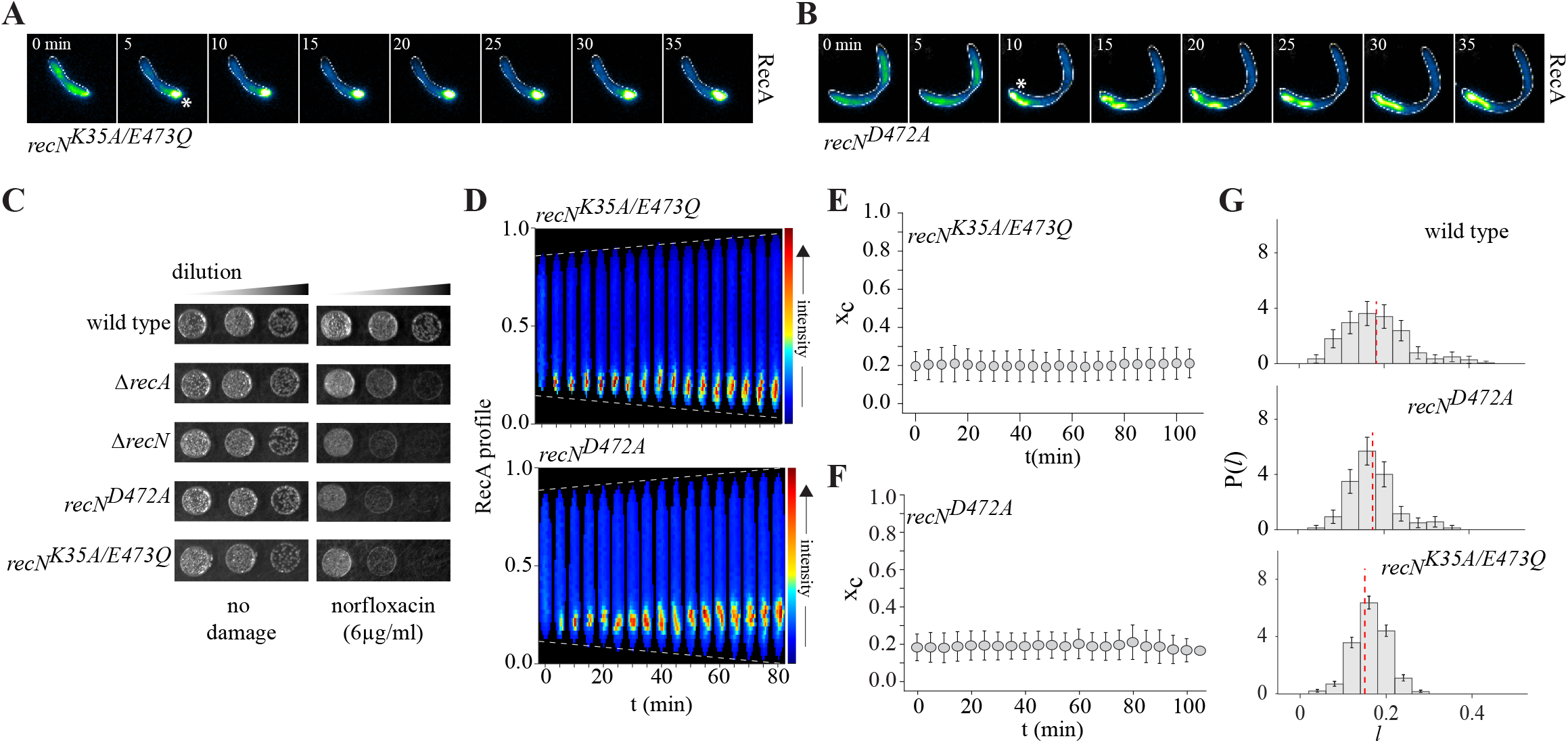
RecN ATPase cycle regulates RecA filament dynamics. (A) Montage of RecA-YFP tracked over time after induction of a single DSB (white asterisk) for putative RecN ATP hydrolysis mutant (*recN^K35A/E473Q^*). (B) Representative montage of RecA-YFP tracked over time after induction of a single DSB (white asterisk) for putative RecN ATP binding mutant (*recN^D472A^*). (C) Sensitivity of *recN^D472A^* and *recN^K35A/E473Q^* to DSB-inducing agent norfloxacin is evaluated (in comparison to wild type, Δ*recN* and Δ*recA* cells, from Fig. S3E). Representative image from at least three independent repeats. (D) [top] Representative RecA-YFP fluorescence profile across the length of the cell in a *recN^K35A/E473Q^* background. [bottom] Representative RecA-YFP fluorescence profile across the length of the cell in a *recN^D472A^* background. (E) Position of centroid of RecA filament normalized to length of the cell (x_*c*_) is plotted for *recN^K35A/E473Q^.* Mean and standard deviation is shown (n = 50). (F) As (E) for *recN^D472A^*. (G) Density histogram of RecA filament lengths (*l*) across all time points of imaging for wild type, *recN^D472A^* and *recN^K35A/E473Q^* (n = 229). Distribution of *l* was found to be significantly different between wild type and *recN^K35A/E473Q^* (p-value = 0.0005, 95% CI [0.008051, 0.02846], Welch’s t-test). Distribution of *l* was comparable between wild type and *recND^472A^* (p-value = 0.1535, 95% CI [-0.003326, 0.02108]). All data shown are from pre-divisional cells.

Taken together, our observations position RecN early in the recombination pathway, specifically in enabling RecA filament translocation during homology search, with influence on RecA dynamics even in the absence of a homologous repair template. The differential impact of putative RecN ATPase mutants on RecA filament lengths (both resulting in abrogation of translocation), leads us to suggest that RecN, via its ATPase cycle, likely modulates RecA nucleoprotein filament remodelling and drives directional translocation of the filament during homology search.

## Discussion

Using a combination of quantitative live cell imaging and theoretical modelling, we have uncovered a new mechanism for RecA action in long distance homology search during recombination, a fundamental process for the maintenance of life. Our first result is that following a site-specific DSB, RecA nucleates at the DSB site and grows into a nucleoprotein filament which moves in a stochastic but directional manner along the length of the cell, in tandem with RecN. The second main result is that the filament translocation is accompanied by dynamic remodelling of the RecA filament. The dynamic remodelling is a consequence of RecN influenced assembly-disassembly that breaks left-right symmetry (akin to treadmilling) - indeed, a stochastic description of this process agrees remarkably well with quantitative analysis of live cell imaging. The RecA filament makes multiple traversals across the length of the cell before the homologous pair is found, followed by repair. That RecN, via its ATPase cycle, is crucially involved in the search, its termination and subsequent repair is the third main result of the paper.

Following our present study, an immediate goal is to deduce the molecular basis of the feed-back regulation that maintains a fixed mean filament length at steady state. Similarly, we will need to unravel the molecular basis of the RecN-regulated dynamics of the RecA nucleoprotein filament. For example, *in vivo* studies in *E. coli* have variably suggested that the DSB-associated RecA structures could either be a bundle^5,58^ or a single^6^ polymer of RecA molecules. It would be informative to determine the detailed composition of the RecA filament in the present study, along with any impact of RecN on the same. This would require a quantitative characterization of the complete RecA-RecN ATPase cycle, its interaction with the mechanochemistry of the DNA substrate, as well as detailed dissection of the RecA filament architecture.

Based on our current observations, we consider two possible scenarios: (A) RecN-mediated regulation of assembly-disassembly rates of RecA molecules in the filament. (B) Motor-driven movement of the RecA nucleoprotein filament and/ or the underlying ssDNA substrate. In line with the possible influence of RecN on mobility of the underlying ssDNA substrate, we find that, in the absence of *recN,* Single-Strand Binding protein (SSB) localization is also restricted to the pole where the break occurs (Fig. S5A). However, pure motor-assisted movement would not be able to explain the stochastic remodelling the filament length, while pure assembly-disassembly, without the dynamics of the DNA, would need to have available a cell-spanning ssDNA segment, which does not appear to be the case in our conditions (Fig. S5A).

Thus, we suggest that the RecN-assisted dynamics of the RecA filament could be a consequence of both motor-assisted movement (such as DNA extrusion) and regulation of assembly-disassembly rates of RecA on DNA (via treadmilling). Fig. S5B is our proposed model of the same. Here, RecN action on the DNA could be a consequence of its reported bridging activity of ssDNA (generated around the break-site) with dsDNA^55^, thus driving directional movement of RecA, along with its underlying DNA substrate. ‘Zipping’ refers to RecN action ahead of the translocating RecA filament, while ‘extrusion’ refers to RecN action behind the filament^72,73^.

Why does the RecA filament need to undergo dynamic remodelling in addition to movement during homology search? Studies have shown that RecA ATPase rates are stimulated 4-fold upon strand invasion, potentially allowing the RecA filament to release from heterologous pairing during microhomology sampling as well^43^. Separately, RecN has been found to (a) stimulate RecA strand invasion activity^49^ and (b) bridge ssDNA and dsDNA^55^. Taken together, it is possible that the RecN-dependent RecA remodelling is a reflection of such microhomology sampling activity prior to homology pairing. The stochasticity of the RecN-driven RecA dynamics and the multiple traversals before identification of the homologous repair template, suggests a novel search mechanism that allows for a regulated genome-wide sampling for homology. Our observations reveal that in addition to the initial waiting time between nucleation to the start of traversals, the translocating RecA filament makes pauses at random intervals. We speculate that RecA molecules might engage in microhomology sampling both initially in the vicinity of the damage site (especially in the post-replicative cohesion period) and during these pause intervals during translocation^43,74^.

Based primarily on *in vitro* observations, earlier models have suggested that RecA engages in 1D sliding movement and intersegment hopping, while regions within the RecA filament engage in microhomology sampling during search^35,74,75^. Our work reveals a central role for regulation of RecA filament dynamics by RecN during long distance search. We uncover the three key elements of this RecN-driven homology search *in vivo*: RecA filament mobility, remodelling of the filament and the ability to undertake multiple pole-to-pole traversals. Together, this suggests a new mechanism of iterative, long distance search within the cell. RecN-mediated multiple pole-to-pole traversals by RecA builds in a kind of *reset* at a fixed spatial location, a strategy that can lead to optimal search^76^, thus facilitating repair well within a single generation time of bacterial growth.

It is possible that there is some diversity in RecA-dependent long-range homology search dynamics across bacterial systems, to contend with the varying cell shapes, chromosome replication dynamics (multi-fork vs replication only once per cell cycle), and chromosome organization^3–6^, thus resulting in static^5,6^ or mobile (present study) RecA structures. However, given that RecA and RecN are among the most conserved proteins of the recombination pathway in bacterial genomes and its counterparts are present across domains of life, we believe that the dynamics we describe here is active in other systems as well.

## Acknowledgements

Authors thank Julia Hitschfel for assistance with bacterial-two-hybrid experiments as well as Meghna Iyer and Varshit Dusad for assistance with developing analysis tools. Authors are grateful to Rodrigo Reyes-Lamothe, David Sherratt, M Srinivasan and AB lab members for feedback and discussion. AB acknowledges support from DST-SERB CRG 2019/003321, HFSP-CDA award (00051/2017) and intramural funding from NCBS-TIFR. JP, SK, KI and MR thank the Simons Foundation for a grant. MR thanks DST-SERB, India for a JC Bose Fellowship.

## Materials and Methods

### Bacterial strains and growth conditions

Strains, plasmids and oligos used in this study are described in Table S1-S3, respectively. Chromosomal modifications such as integration of fluorescent markers or deletion of genes were performed using the two-step recombination method^77^ or with vectors described previously^78^. Transductions were carried out using ΦCr30^79^. Unless otherwise stated, *Caulobacter* cultures were grown at 30°C in peptone yeast extract and supplemented with antibiotics at appropriate concentrations. When growing strains with *dnaA* under an IPTG-inducible promoter, liquid media was supplemented with 0.5 mM IPTG and solid media with 1 mM IPTG. For synchronization experiments, *Caulobacter* cultures were grown till mid-log phase and synchronization protocols were followed as previously described^4^. Protocols used for isolating non-replicating pre-divisional cells and non-replicating swarmer cells^4,38^ are briefly described here. For isolation of non-replicating swarmer cells, cultures were grown till mid-log phase in media supplemented with IPTG. Cells were then depleted for DnaA by washing off the inducer and were grown in the absence of IPTG for one generation. This was followed by synchronization and isolation of swarmer cells. These cells were re-suspended in media without IPTG and used for DSB induction experiments. For isolation of non-replicating pre-divisional cells, cultures were grown till mid-log phase in media supplemented with IPTG. Cells were then synchronized and swarmer cells isolated. Swarmers were then released into media lacking IPTG. Cells were then depleted for DnaA by washing off the inducer and grown in the absence of IPTG for one generation. Cell division was blocked via addition of 35 μg/ml cephalexin and these non-replicating pre-divisional cells were further used for DSB induction experiments. In all cases, *recA-YFP* was induced using 0.03% xylose 90 min prior to imaging and subsequently maintained in the agarose pad as well during the entire course of imaging. For imaging SSB-YFP after induction of a break in non-replicating swarmer cells, *ssb-YFP* was induced using 0.03% xylose. DSBs were induced with 2 μM Vanillate. For this, vanillate was added to the growth culture 15 min prior to imaging and was subsequently maintained on the agarose pad as well.

### Fluorescence Microscopy

Time course imaging was performed on 1% agarose pads (Invitrogen ultrapure). For time-lapse imaging, cells were grown on 1.5% GTG agarose pads (low melting) prepared in peptone yeast extract and supplemented with vanillate, xylose or cephalexin at appropriate concentrations and imaged using glass-bottom petri dishes. Imaging was carried out using a wide-field epifluorescence microscope (Eclipse Ti-2E, Nikon) with a 63X oil immersion objective (plan apochromat objective with NA 1.41), illumination from pE4000 light source (CoolLED), Hamamatsu Orca Flash 4.0 camera and a motorized XY stage. During time-lapse imaging, focus was maintained using an infrared-based Perfect Focusing System (Nikon). Image acquisitions were done using NIS-elements software (version 5.1) and images were acquired every 30s, 1 min or 5 min (as indicated in the main text and respective figure legends). For most experiments with RecA, exposure time used for excitation at 490 nm was 400 ms and for 550 nm was 500 ms. For imaging MipZ, exposure time used for excitation at 550 nm was 100 ms. For imaging YFP expressed from the *sidA* promoter, exposure time used for excitation at 490 nm was 500 ms. In all images, scale bars = 2μm. Procedures for image analysis and for extracting key features of RecA localization are described in *Supplementary Results.*

### DNA degradation assay

Implemented as previously described^38^. Briefly, non-replicating swarmer cells with DSB site at +780kb were isolated and released in a media with 0.5 mM Vanillate to induce DSBs. Samples were collected at 0 h (control) and 1 h after DSB induction. Genomic DNA was isolated using DNAeasy blood and tissue kit (Qiagen). Whole genome Illumina sequencing was carried out on the isolated DNA (NGS facility, NCBS, India). Data were further analysed using the following protocol. First, indexing with the reference genome (4.01 Mbp) (NCBI reference sequence:NC-011916.1) was done using BWA^80^. Reads with raw read quality 20 were aligned using ‘BWAaln −q’. SAMTOOLS (version 0.1.19-96b5f2294a)^80^ was used to filter out the multiply-mapped reads. Finally, with BEDTOOLS^81^ the read count per bin was calculated using the .bed files containing bin positions. *Caulobacter* genome data obtained after Illumina sequencing, Hiseq 2500 illumina short reads (50 bp), was divided into 1 kb bins. Read counts per bin were normalized to the total reads acquired for that sample. The ratio of normalized reads after DSB (1 h) to normalized reads before DSB (0 h) was plotted to visualize the read enrichment profile obtained across the genome after induction of a DSB. The graph generated was processed further with the Lowess smoothing function in MatLab.

### Bacterial two-hybrid assay

Implemented as previously described^38^. Briefly, to investigate physical interaction between a pair of proteins, their respective genes were fused to 3’ end of T25 or T18 fragments in pKT or pUT vectors. These vectors were co-transformed into *E. coli* BTH101. Co-transformed cells were grown to saturation in M63 media with maltose and IPTG and 5μl of this culture was spotted on MacConkey agar plates (40 g/L) with maltose, IPTG and appropriate antibiotics. Plates were incubated at 30°C for 2-3 days.

## Supplementary content

Supplementary Results with detailed description of theory and quantitative analysis. Supplementary figures (associated with main text).

Supplementary videos 1-6. Supplementary tables 1-3.

## Supplementary material

### Supplementary Figures

**Fig. S1:**
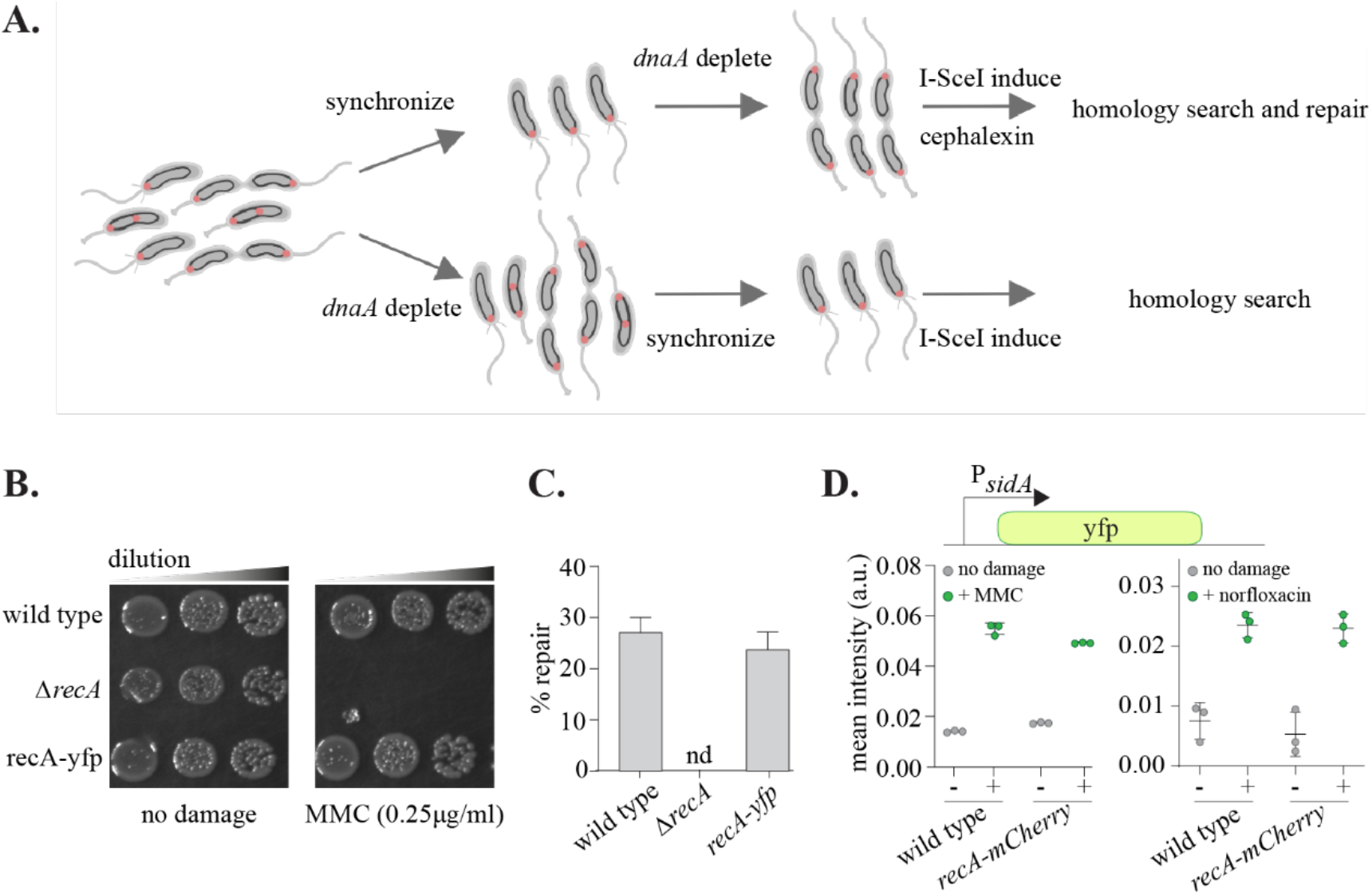
Dynamics of RecA filament during homologous recombination. **(A).** Schematic of experimental setup. *Caulobacter* cells carrying either two completely replicated and segregated chromosomes or a single non-replicating chromosome (black circles inside the cell) are isolated as previously described. DSB site is integrated on the chromosome near the origin of replication and visualized via fluorescently-tagged MipZ (red dot), that forms foci at the polarly localized origin regions in *Caulobacter.* DSB is induced via regulated expression of a destabilized I-SceI enzyme. Cells with 2n chromosome content (pre-divisional cells) will carry out all steps of recombination (from homology search to repair) while cells with 1n chromosome content (swarmer cells) can be assessed for their ability to carry out homology search alone. **(B).** Strain expressing *recA-yfp* does not show increased sensitivity to generic DNA damaging agent, mitomycin-C (MMC), when compared with wild type cells (representative image from 3 independent repeats). **(C).** Repair events (reappearance of both MipZ localizations after loss of a single localization) are assessed across a 2 hr window of imaging for all cells that experienced a single DSB (as measured by loss of one but not both MipZ localizations). % cells showing repair is similar to previously published reports. No repair is detected (nd) in cells lacking *recA* (mean and standard deviation from three independent repeats (n= 100 cells for each repeat)) **(D).** RecA-YFP expressing cells are SOS proficient with similar induction levels as seen in the case of wild type cells. SOS induction levels are estimated by assessing the activity of the *sidA* promoter expressing a fluorophore. Cells are treated with MMC or Norfloxacin for 2 hrs. Mean intensity is plotted for three independent repeats (n= 200 cells for each repeat).

**Fig. S2:**
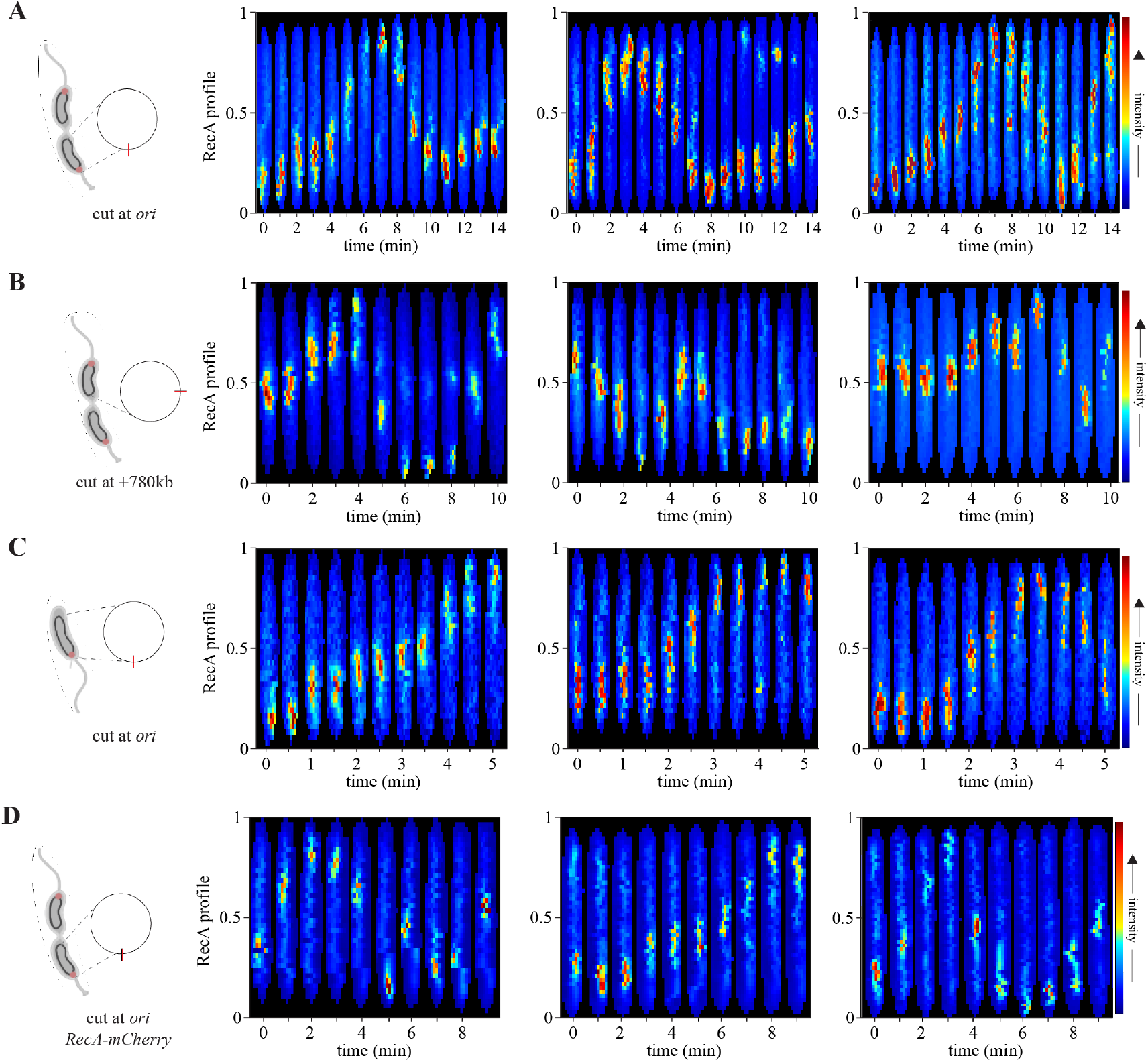
Directional and dynamic movement of RecA filament during homology search. **(A).** Representative RecA-YFP fluorescence profiles across the length of the cell over pole-to-pole traversal events during homology search. DSB is induced near the origin of replication in non-replicating pre-divisional cells with two fully replicated and segregated copies of the chromosome. In all panels in this figure, *t0* is defined as the time point prior to the observation of a positive displacement of the RecA localization from the pole at which it first localized (DSB pole) (**B).** As (A) for DSB induced +780kb from the origin of replication. In this case, localization starts near ¼ or ½ position and proceeds towards either cell pole. **(C).** Representative RecA-YFP fluorescence profiles across the length of the cell over pole-to-pole traversal events during homology search. DSB is induced near the origin of replication in nonreplicating swarmer cells with a single chromosome, thus lacking a homologous template for repair. **(D).** Representative RecA-mCherry fluorescence profile across the length of the cell show comparable dynamics to that observed in case of RecA-YFP in (A).

**Fig. S3:**
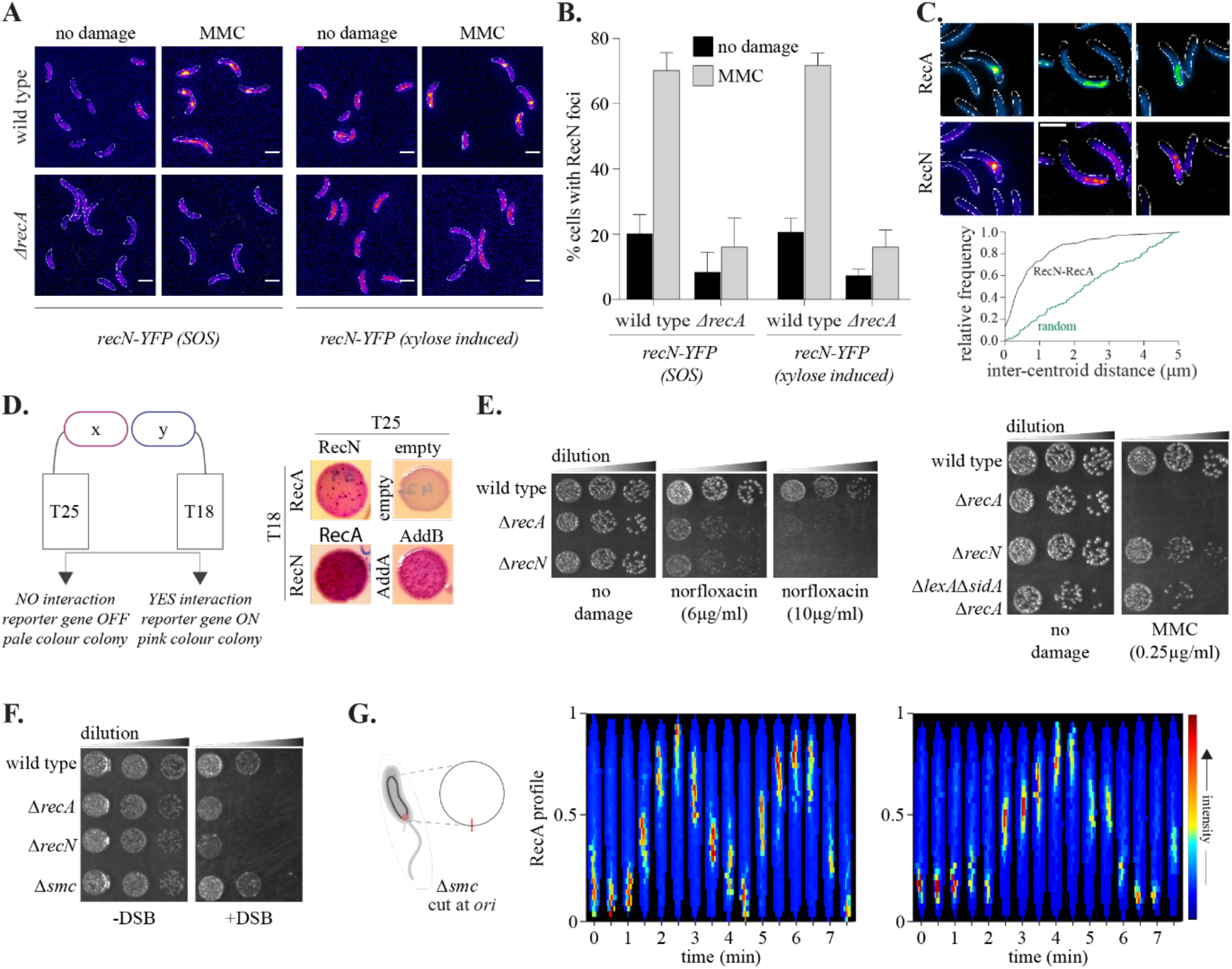
RecN is essential for RecA filament movement *in vivo.* **(A-B).** RecN localization under DNA damage (MMC) is dependent on RecA. To distinguish between SOS induction requirement or RecA requirement, *recN* is also expressed from a xylose inducible promoter. In this case as well, localization in response to damage is dependent on presence of RecA (*n* = 100 cells for each condition, mean and standard deviation from three independent repeats). **(C).** RecA and RecN colocalize at a DSB. Representative images of RecA-mCherry and RecN-YFP are shown, after induction of a DSB at *ori.* Below, distance from centroid of RecN localization to centroid of RecA localization is measured (*n* = 100 localizations). As a control, distance from centroid of RecN to any pixel within the cell is plotted (random). **(D).** Physical interaction between RecN and RecA is assessed using the bacterial-two-hybrid assay. RecA/ RecN are fused to the T25 or T18 fragments of the adenylate cyclase gene. Positive interaction would bring the two fragments together and result in the expression of a reporter gene from a cAMP-inducible promoter, resulting in production of pink/ red colonies. Lack of interaction would result in pale colony colour. As a positive control AddA and AddB interaction is tested. *‘empty’* refers to the T25 and T18 fragments alone. Representative image from at least three independent repeats. **(E).** *ΔrecA* or *ΔrecN* cells are similarly sensitive to DSB-inducing agent norfloxacin. In case of MMC, which causes generic DNA damage, *ΔrecA* cells show higher sensitivity than *ΔrecN.* However, cells lacking *recA* with constitutive SOS response ON (*ΔlexAΔsidAΔrecA*) have similar sensitivity to MMC as *ΔrecN,* suggesting that both proteins are similarly required for MMC damage repair (excluding the function of RecA in regulating the SOS response). Representative image from at least three independent repeats. **(F).** *ΔrecA* or *ΔrecN* cells are similarly sensitive to DSB induction via I-SceI. In contrast, *Δsmc* cells show comparable growth to wild type cells. Representative image from at least three independent repeats. **(G).** Representative RecA-YFP fluorescence profiles across the length of the cell in cells lacking *smc.* Here, *t0* is defined as the time point prior to the observation of a positive displacement of the RecA localization from the pole at which it first localized (DSB pole).

**Fig. S4:**
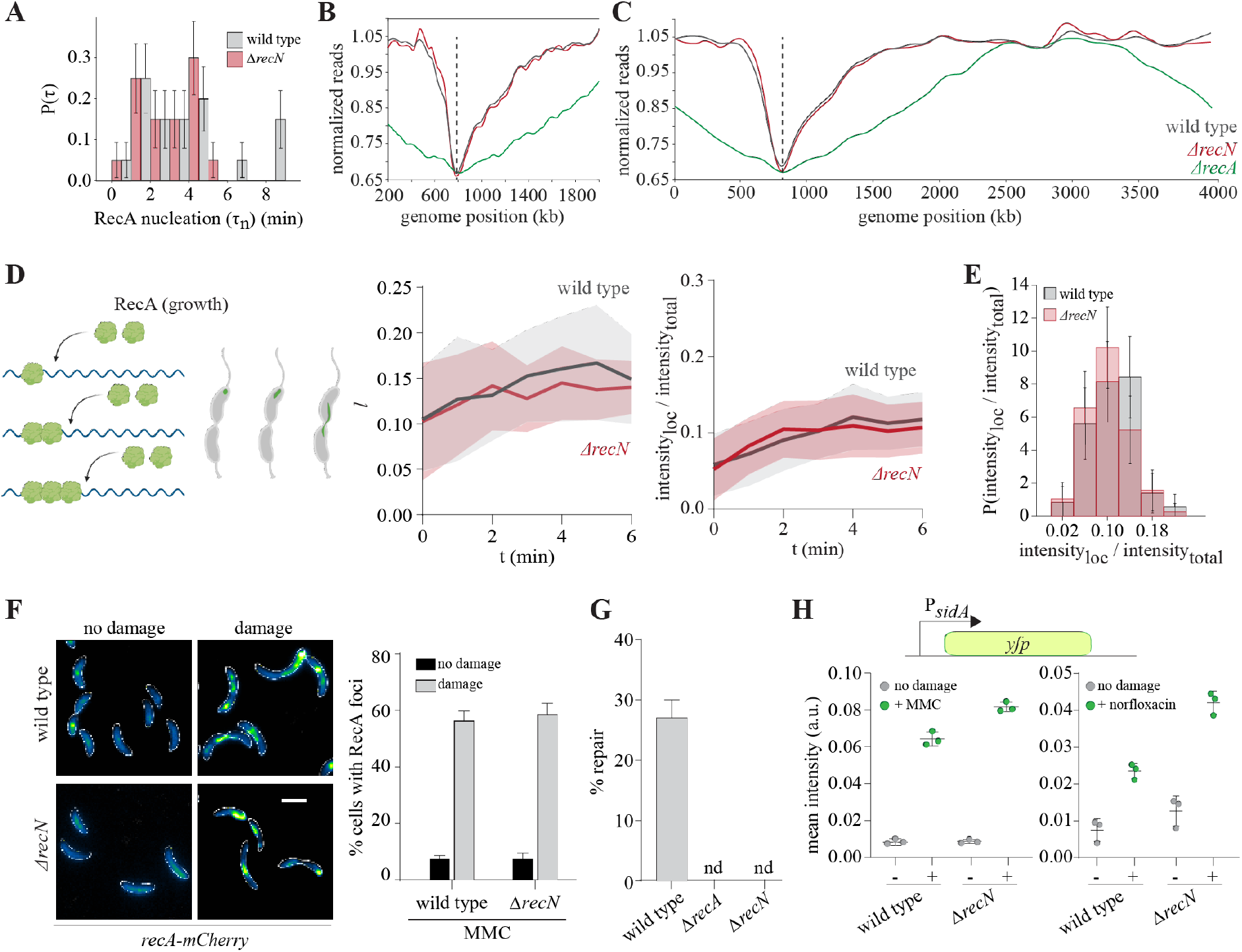
RecN does not impact RecA nucleation. **(A).** Probability distribution of RecA nucleation time (time between disappearance of a single MipZ marker (*t0*) and appearance of RecA localization at that position) for wild type (grey) and *ΔrecN* (red) cells (*n* = 100). **(B)**. Zoomed in (200 -2000 kb) DSB processing profiles for cells induced with break at +780kb (dashed line) in wild type (grey), *ΔrecN* (green) and *ΔrecA* (red) cells. Sequencing reads are normalized to a no damage control and loss of reads around the break site is indicative of loss of double-stranded DNA. Cells lacking *recA* undergo ‘rec-less’ degradation, resulting in significant loss of sequencing reads around the DSB site. Representative plot from two independent repeats. **(C)** Whole genome profile for data in **(B)**. **(D).** RecA filament growth (time between RecA nucleation (*t0*) and start of first translocation cycle (in case of wild type)) is measured either as a function of change in RecA filament length (*l*) over time [right] or change in localization intensity over time (intensity_*loc*_/ intensity_*total*_) [left] for wild type and *ΔrecN* cells. Solid line represents the mean and shaded region the standard deviation. **(E)** For reference, distribution of localization intensities for all cells is also shown (n=50). **(F).** RecA localization is dependent on DNA damage. Percentage cells with RecA localization with or without MMC damage is plotted for wild type and *ΔrecN* cells (mean and standard deviation from three independent repeats, *n* = 100). **(G).** I-SceI-induced DSB repair events are estimated for wild type, *ΔrecA* and *ΔrecN* cells. Repair in absence of *ΔrecA* or *ΔrecN* is not detectable (nd) in a 2 hr window of imaging (n=100 cells for each repeat). Wild type and *ΔrecA* measurements are from Fig. S1C (mean and standard deviation from three independent repeats). **(H).** SOS induction levels are measured via P_*sid4*_-yfp expression under MMC or Norfloxacin damage for wild type and *ΔrecN* cells (mean and standard deviation from three independent repeats (n=200 cells for each repeat)).

**Fig. S5:**
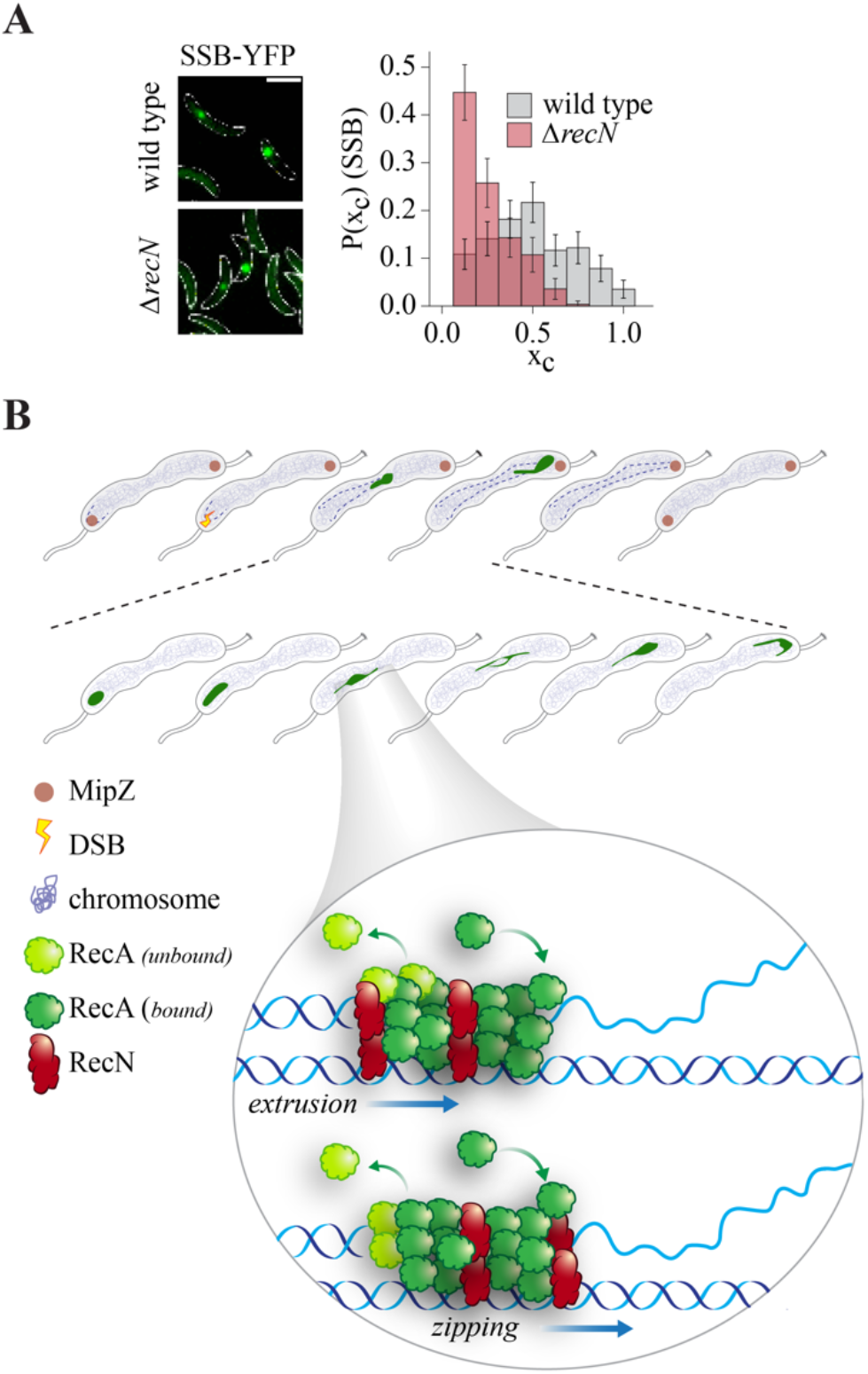
Proposed mechanism for RecN-driven RecA filament translocation and remodeling during homology search. **(A).** [Above] Representative images of SSB-YFP localization upon induction of a single DSB in non-replicating swarmer cells. [Below] Probability distribution of SSB-YFP centroid position (x_*c*_) in wild type (*n* = 369) and *ΔrecN* cells (*n* = 278). **(B).** Proposed mechanism for RecN action on the RecA filament. Upon induction of a DSB near the polarly-localized origin, we observe localization of RecA to the break site. The RecA localization then dynamically translocates across the length of the cell undergoing several pole-to-pole traversal events before the homologous partner is found. This movement and associated remodeling of the nucleoprotein filament depends on RecN. We propose that RecN modulates RecA filament lengths during translocation via regulating rates of RecA subunit assembly-disassembly. In addition, RecN action on DNA (via possible bridging activity between ssDNA and dsDNA) can facilitate RecN-driven translocation of the RecA filament and its underlying DNA substrate. We use ‘zipping’ to refer to RecN action ahead of the filament and ‘extrusion’ for action behind the filament.

### Supplementary Video Legends

Video 1: RecA-YFP in pre-divisional cells after induction of a single DSB near the polar origin of replication. RecA-green; cell outline-white. Four individual cells shown. Time in min.

Video 2: RecA-YFP in swarmer cells after induction of a single DSB near the polar origin of replication. RecA-green; cell outline-white. Four individual cells shown. Time in min.

Video 3: RecA-YFP in pre-divisional cells lacking *recN,* after induction of a single DSB near the polar origin of replication. RecA-green; cell outline-white. Four individual cells shown. Time in min.

Video 4: RecA-YFP in swarmer cells lacking *recN,* after induction of a single DSB near the polar origin of replication. RecA-green; cell outline-white. Four individual cells shown. Time in min.

Video 5: RecA-YFP in pre-divisional cells carrying *recN^D473A^* mutant, after induction of a single DSB near the polar origin of replication. RecA-green; cell outline-white. Four individual cells shown. Time in min.

Video 6: RecA-YFP in pre-divisional cells carrying *recN^K35A/E473Q^* mutant, after induction of a single DSB near the polar origin of replication. RecA-green; cell outline-white. Four individual cells shown. Time in min.

## Supplementary Results

### S1. QUANTITATIVE ANALYSIS OF EXPERIMENTAL DATA

Images were analysed using Microbetracker/ Oufti [1, 2] and MatLab or manually assessed in ImageJ. In all cases, experiments were biologically replicated at least three times and pooled data from at least three independent repeats are plotted (after confirming reproducibility between replicates). Plots were generated using custom written code, MatLab (2019, 2020), R, Python or GraphPad Prism 8.0. Schematics were made using Biorender and Adobe Illustrator. Following image acquisition, phase contrast profiles of cells were segmented using Oufti. Mesh files were then overlayed on the raw images (.tiff). ‘roipoly’ function in Matlab was implemented to extract pixel values only within the desired cell ROI. Given the curved nature of Caulobacter cells, for the ease of analysis the ROIs of segmented cells were straightened. For this, firstly the ROI was rotated by angle equal to the angle between the major axis of the ROI and x-axis. This angle was estimated by extracting the image property ‘Orientation’ using MatLab function ‘regionprops’. Secondly, a line was drawn across the cell length, positioned at mid-cell. This was taken as the reference line to which all pixels of the ROI were re-centered. Following this straightening, pixels were adjusted along the mid-cell line to maintain all cell information (cell area and fluorescence).

RecA nucleoprotein filament properties were analysed via two key dynamical variables - position of the centroid of RecA filament relative to the length of the cell (x_*c*_) and length of the RecA filament relative to the cell length (*l*). In addition, intensity (*i*) was also estimated, as the ratio of RecA fluorescence within the localization to total cell fluorescence. To obtain the above-described parameters, an eCDF (Empirical Cumulative Distribution Function) plot was generated for the fluorescence intensities per pixel in every ROI throughout the time-lapse movie. From this, intensity value ? 96% of the eCDF plot was selected from a single frame as the threshold for automated detection of RecA nucleoprotein filament localizations, that allowed for accurate detections of most localizations in the movie. This threshold was then uniformly applied across for all frames of the time-lapse. Values below this threshold were considered as part of background fluorescence coming from the freely-diffusing pool of RecA-YFP and were not included for analysis of (x_*c*_) or (l). RecA fluorescence profiles thus generated were visualized in the form of a kymograph generated in MatLab. Profiles were compared to the raw images obtained to ensure that false positive or false negative detections were avoided.

Δx_*c*_ and Δ*l* were calculated by subtracting the value at the present frame from the value at the previous frame. To estimate velocity v_*c*_, a least square fit of degree 1 was made for the x_*c*_ values during a traversal using ‘polyfit’ function in python. The slope of this linear fit was defined as the mean velocity of a traversal, v_*c*_. The v_*c*_ values falling above or below the range 0.006 - 0.001 were considered as outliers, traversals with only 2 data points were excluded. These features were analyzed for cells during the ‘homology search’ phase as defined by the first recordable positive non-zero value for Δx_*c*_, following the nucleation and growth of the RecA localization (Fig. S1A). The first time point x_*c*_(0) was set to the frame before the Δx_*c*_ value was non-zero. The last time point for a single traversal was recorded when the centroid of RecA localized at the opposite cell pole from where the mobility had started. We estimate that this phase represents a steady-state dynamic based on the following observations: a. distribution of *l* in invariant across time-points of imaging during the homology search phase (Fig. S1C-E). b. distribution of *l* is comparable between pre-divisional and swarmer cell states (where homology search is continuous, due to the inability to complete recombination-based repair) (Fig. S1B).

The nonlinear fits to the density histograms of (*l*) in the cases of wild type and *recN* deleted cells were generated using in-built functions in Mathematica. Using the respective histogram data, parameters in the equations (32) and (48) were estimated using the function “NonlinearModelFit” which uses the Levenberg-Marquardt algorithm. Reasonable initial conditions needed to be provided to the function/algorithm in order to ensure plausible estimates of the parameters. With these estimates, consistency checks for the probability density functions were done and the curves/graphs (green) plotted (Fig. S8-S9).

The > 95% confidence intervals to calculate Margin of Error (MoE) for probability distributions were estimated according to the formula for population proportion estimator,

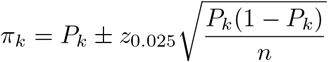

where *n* is the data/sample size, *P_k_* is the empirical proportion calculated using the data for bin *k,* and *π_k_* is the true population proportion for the bin *k*. To estimate the density, however, the above equation was divided by the respective bin widths *δx_k_*.

**FIG. S1.**
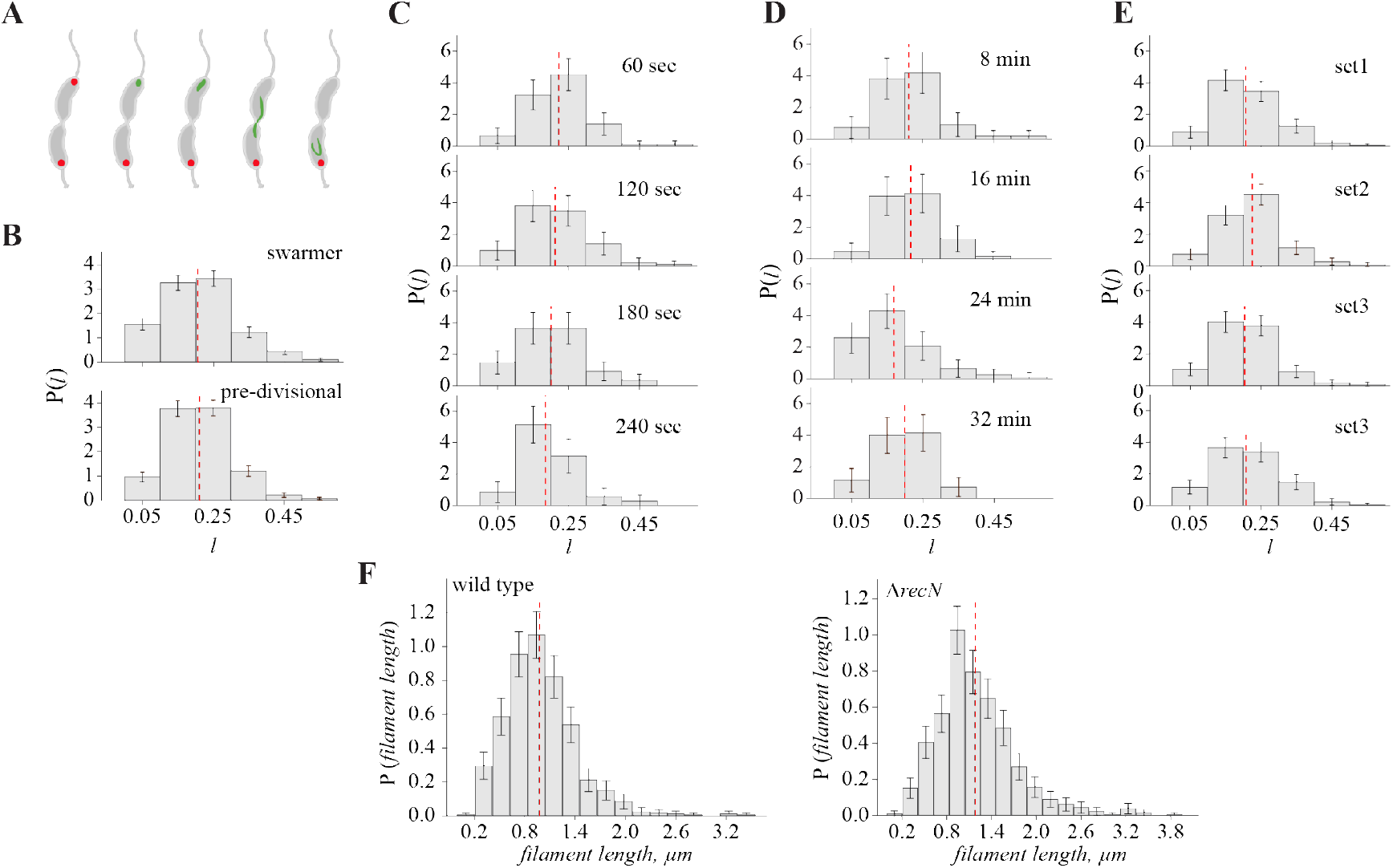
A. Schematic depicting steps of RecA nucleation, growth and homology search. B. Density histogram of RecA filament lengths during homology search in swarmer and pre-divisional cells indicates that lengths are comparable (*n* = 858). C. Density histogram of RecA filament lengths at indicated time intervals during homology search indicates that lengths are comparable. Same set of cells are tracked across all time points (*n* ≥ 70). D. Density histogram of RecA filament lengths during homology at indicated time windows of imaging indicates that lengths are comparable (*n* ≥ 70). E. Density histogram of RecA filament lengths for subset of cells (25 cells in each set) from length distribution in Fig. 3E. F (*n* ≥ 214). Density histogram of RecA filament lengths (in *μ*m) for wild type (*n* = 795) or *recN* (*n* = 852) deletion mutants. Fraction length (*l*) is calculated from these values. In all plots, red dashed line is indicative of mean.

### S2. STOCHASTIC DESCRIPTION OF THE RecN-ASSISTED DYNAMICS OF THE RecA FILAMENT

We provide a quantitative stochastic description of the mechanochemistry of RecN-assisted RecA filament dynamics.

#### A. Mechanochemistry and Assembly-Disassembly Cycle

##### 1. Mechanochemical states

To do this, we first describe the conformation states of the RecA-RecN system, both unbound and bound to the DNA strand. RecA and RecN are both ATPases, and hence should exist in their native states and in their ATP-bound and ADP-bound states [3–7].

From [8–10] we know that RecA-ATP is bound to the ssDNA, while RecA-ADP is primed for unbinding from the DNA substrate. While there is no information yet on the DNA binding status of RecN-ATP or RecN-ADP, we do know that a form of RecN binds to the DNA bound RecA-ATP [6]. Thus we will simply assume that there is, in addition to the native RecN, a bound state of RecN.

This is the conformation state space of the RecA-RecN system that we display in Fig. S2.

**FIG. S2.**
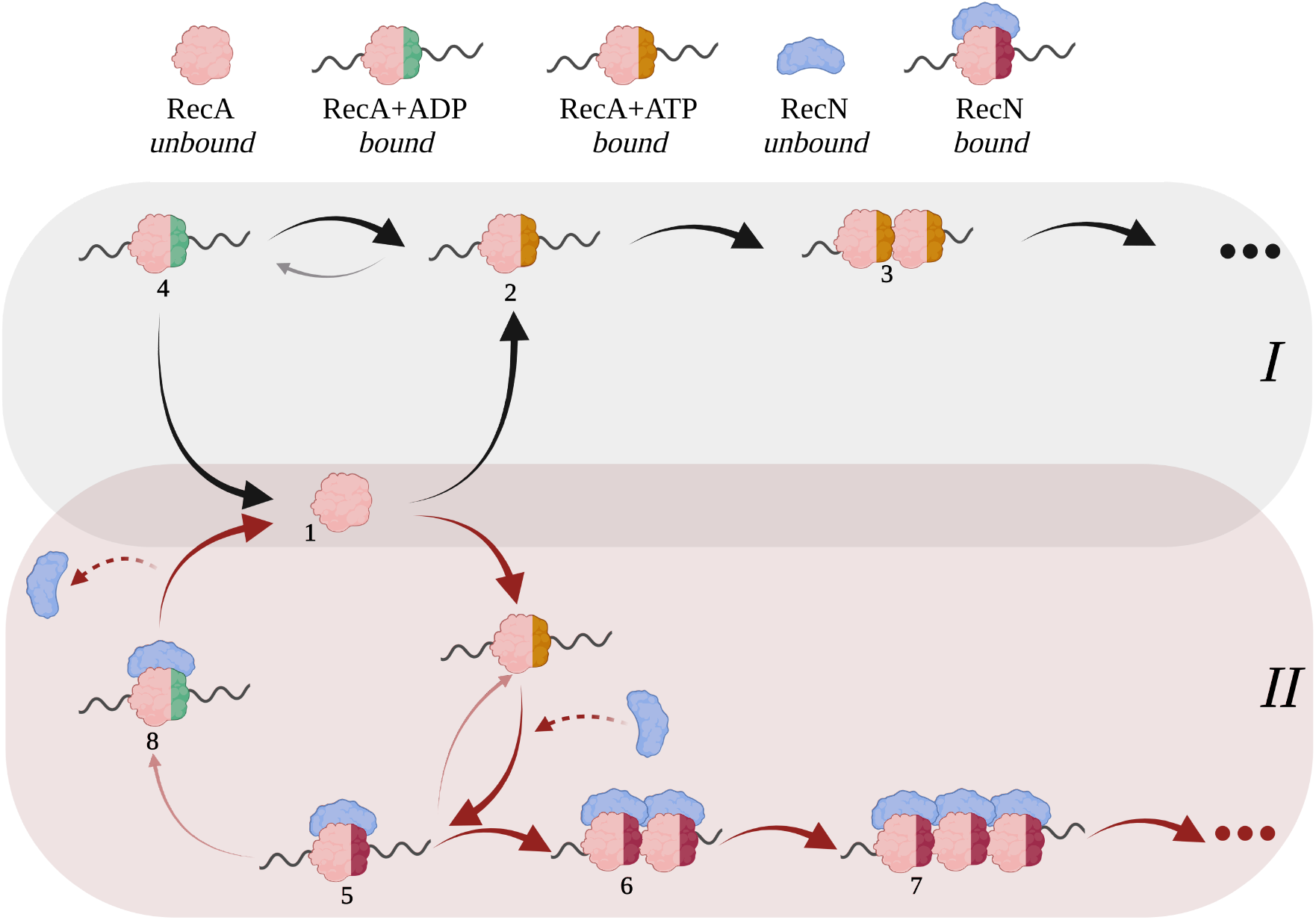
(top) The conformation states of RecA and RecN, depend on their ATPase activity and their binding to the DNA substrate. (bottom) The assembly-disassembly network of RecA-RecN comprises two modules I and II, described in the text. Thick (thin) arrows denote higher (lower) rates. The directional bias in both the RecN-assisted RecA filament movement (in Module II), and RecA assembly in the recN mutant (Module I), is schematised here by a structural asymmetry of the bound states, which ultimately result in a left-right asymmetry in the binding-unbinding rates onto the DNA strand.

##### 2. Assembly-Disassembly Cycle

We now describe the assembly-disassembly network of RecA and RecN on the DNA strand. This network comprises two strongly connected modules - Module I, which involves RecA alone, and Module II, which involves both RecA and RecN.

The Module I network is displayed in Fig. S3. Here the unbound RecA (node 1) can assemble on the DNA in the RecA-ATP bound state (node 2). From node 2, the RecA-ATP can either transform to RecA-ADP (node 4) by hydrolysis or can assemble an additional RecA-ATP to form a dimer (node 3). From node 3, the RecA-ATP dimer can either transform to node 5 by hydrolysis or can assemble an additional RecA-ATP to form a trimer (node 6), an so on (represented by •••). The polymerization happens unidirectionally along the DNA strand, from the *5’* to the *3’* end.

In Fig. S3, the thick black arrows denote transitions with higher rates, whereas the thin gray arrows denote transitions where the rates are low. It follows that in the absence of RecN, the RecA monomers can in principle be quickly assembled on the DNA substrate from the 5’ to the 3’ end to form a growing RecA filament.

**FIG. S3.**
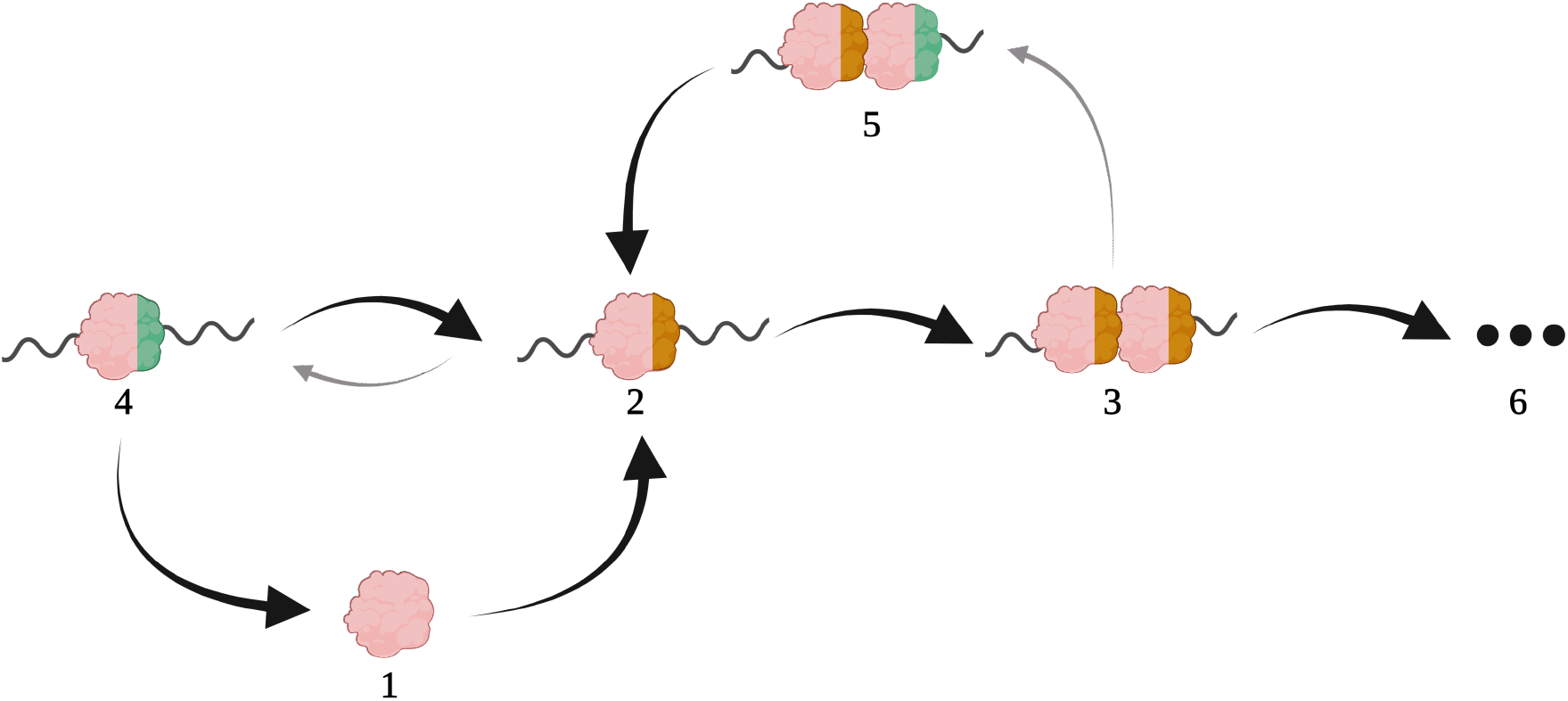
Module I shows the assembly-disassembly network of RecA on DNA. The chemical reaction network comprises a set of nodes that transition from one to another via directed and weighted edges (arrows) that denote reaction paths. The darker arrows are associated with higher reaction rates.

Module II involves both RecA and RecN, and is shown in Fig. S4. Here the unbound RecA (node 1) can assemble on the DNA in the RecA-ATP bound state (node 2), which with the availability of unbound RecN gets transformed to a bound RecA-RecN complex (node 3). From node 3, the bound RecA-RecN complex can either hydrolyse to form RecA-ADP bound to RecN (node 6), followed by disassembly to node 1, or can assemble an additional RecA-RecN complex to form a dimer (node 4), and so on, ……, to form a filament of the RecA-RecN complex. As before, the thick red arrows denote higher transition rates, whereas the thin pink arrows denote transitions where the rates are low.

**FIG. S4.**
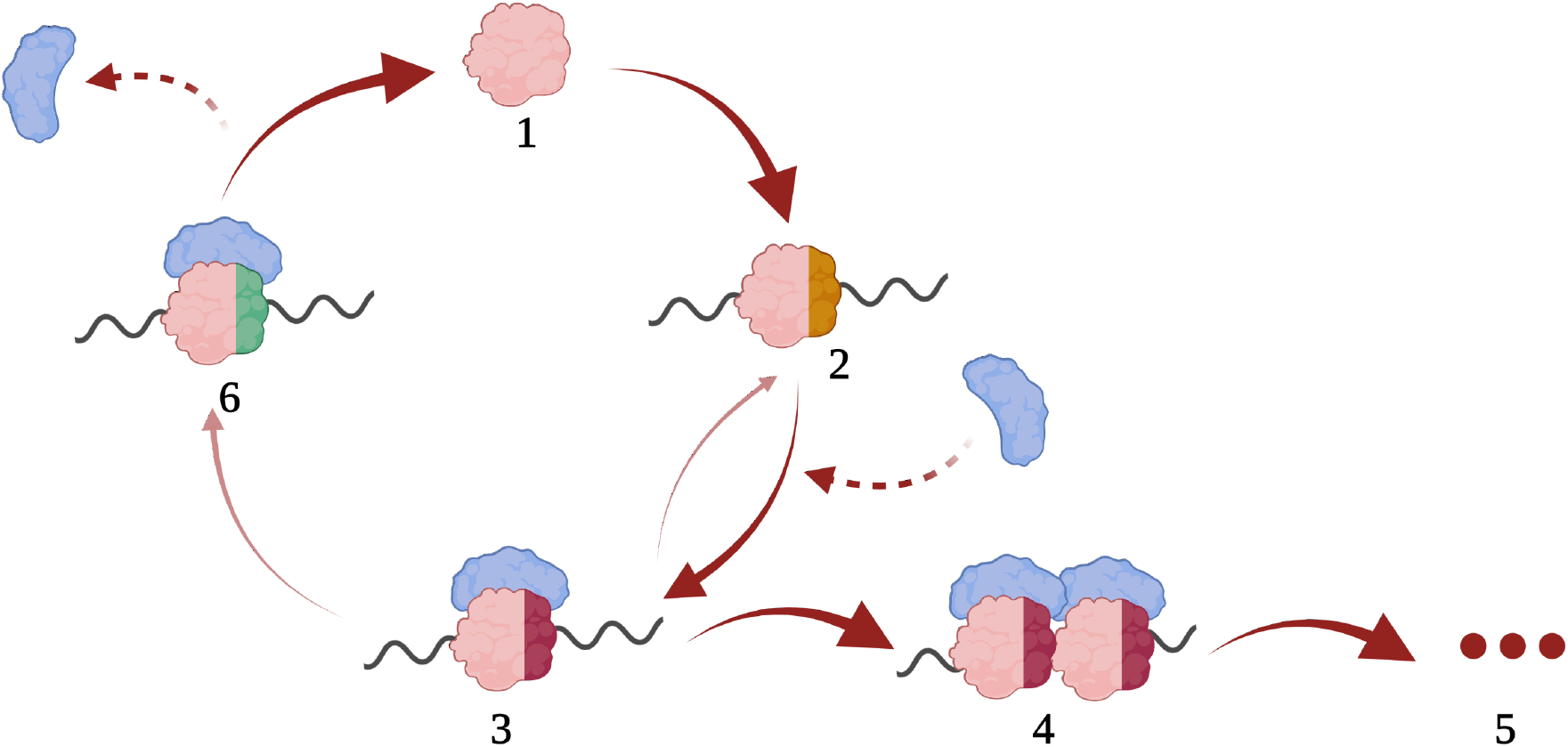
Module II shows the assembly-disassembly network of RecA-RecN on DNA. The blue crescent protein represents unbound-RecN, which enters and leaves this module via the dashed arrows. Here again, the darker arrows are associated with higher reaction rates.

##### 3. Reduction to an effective model

The full network depicted in Fig. S2, of which Modules I and II are a part, is complex and analytically intractable. We therefore need to come up with a suitable reduction of the network which has fewer nodes and directed edges, while retaining the relevant features of the original. Here we present a systematic procedure to do so.

Our strategy is to prune those reactions in the network that do not contribute significantly to network output. Our method is based on a reaction flux analysis of stochastic chemical reaction networks [11–13], which monitors the intensity of the flow between chemical species over a given time interval. We then identify reactions that have low mean reaction flux, compared to a threshold, and eliminate the associated vertices and directed edges, together with modifying the edges in the reduced network. To implement this, we choose a node *i* and compute the signed vertex flux at that node, *F_i_* = ∑_*l_i_*_ *F_i,l_i__*, where the sum is over all directed edges that enter or leave node i. If *F_i_* < *F_min_*, a threshold, the *i*-th node will be removed, except for the case where the *i*-th node represents a decisive reactant. All edges that were connected to at least one of the removed nodes will be removed, too. All other reactions are retained.

This coarse-graining strategy works best for a chemical network with a sequence of reactions that occur in series, with offshoots that are in parallel [11–13], which is precisely the network architecture displayed in Modules I and II.

Applying this procedure to the reaction network in Module I (Fig. S3), we see that node 4 has two strong arrows going away from it, and so the vertex flux of node 4 is very low. We thus prune this part of the network and introduce a directed edge from node 2 to node 1. Similarly node 5 can be eliminated, followed by introducing a directed edge from node 3 to node 2. This procedure can be carried out across the network. The rest of the nodes (and edges) such as 1, 2, 3, are retained. This leads to the reduced Module I shown in Fig. S5.

We prune Module II in the same way. Thus node 6 can be eliminated, whilst introducing a directed edge from node 3 to node 1. Likewise, node 2 can be eliminated, with the introduction of a strong directed edge from node 1 to node 3. The RecN that enters the reaction module with a dashed edge are decisive reactants and will be retained. These considerations lead to the reduced Module II shown in Fig. S5.

It is this reduced ladder model that is amenable to analytical treatment and will be investigated further. Before doing that, there are two comments that we would like to make regarding the reduced network. One, is that since RecA and RecN are ATPases, some of the directed edges in the network are driven by energy transduction and hence violate the condition of detailed balance. The second, is that both the systematic polymerisation of RecA in Module I and the RecA-RecN complex in Module II that is observed in the ΔrecN and wild type cells, is unidirectional from the 5 to 3 on the DNA strand. This breaking of left-right symmetry (parity), must have a manifestation in a structural feature of both DNA-bound RecA, and DNA-bound RecA-RecN complex, that has been schematised in Figs. S3-S5.

**FIG. S5.**
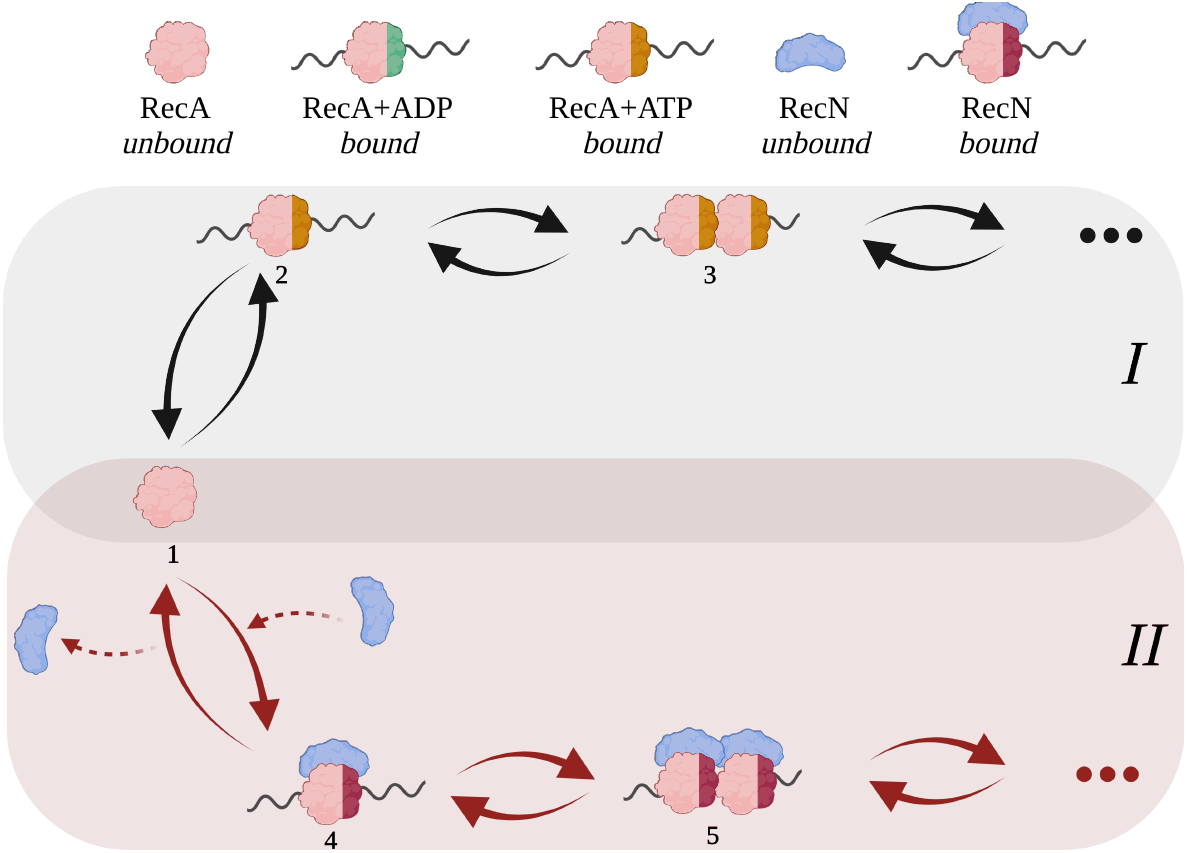
Reduction of RecA-RecN assembly-disassembly following a Flux-based analysis. Here we prune the original network to reduce the number of vertices and directed edges without significantly changing the network flux.

#### B. Dependence of assembly-disassembly rates on *l*

To specify the reduced reaction network completely, we need to provide the assembly-disassembly rates associated with the directed edges. Ideally this should be obtained from independent experiments, such as FRAP, however this poses quite a challenge. Instead we extract some information about these microscopic rates from the experiments described in Figs. 1, 2 and 3 of the main manuscript.

We first note that the distribution of the centroid position x_*c*_ is uniform across the cell (Fig. 1E [inset]) and that the joint distribution P(x_*c*_, *l*) in Fig. 3C reveals that there are no “hot-spots”. This would suggest that the dynamics of *l* does not depend on x_*c*_. Further, the distribution of step sizes, both Δx_*c*_ and Δ*l* show positive and negative values, suggesting that the dynamics of the filament proceeds via assembly and disassembly as it traverses across the cell (Fig. 1E and Fig. 3A).

The most important aspect of the experimental data is that the distribution of filament length P(*l*) at steady state shows a peak at a finite value of *l* = *l*\*, a value that is significantly smaller that the cell length in both the wild type and the Δ*recN* cells (also the filament never extends over the cell length). This we argue, implies that the rates of assembly and disassembly cannot be constant. For, if the assembly rate is larger than the disassembly rate, then the filament will grow without limit. With the same token, if the disassembly rate is larger than the assembly rate, then the filament will shrink to zero. The dynamical flows corresponding to these scenarios is shown in Fig. S6 (i), (ii). The constant assembly-disassembly rates cannot be made to balance each other robustly at a particular length and will not give rise to the observed finite peak *l*\* at steady state.

This implies that the assembly-disassembly rates must depend on the length of the filament. Clearly, to obtain a peaked distribution at *l*\*, one should arrange that assembly exceeds disassembly at *l* < *l*,* and disassembly exceeds assembly at *l* > *l*\*. Such a *negative feedback* between the assembly-disassembly rates and the filament length Fig. S6 (iii), (v), provides a homeostatic balance leading to a finite mean filament length with small fluctuations about it. This results in a stable fixed point at *l* = *l*\* in the flow diagram Fig. S6 (iii), (iv).

It is clear that if the feedback is positive, then the filament will either show unbridled growth or complete shrinkage resulting in an unstable fixed point *l*\*.

**FIG. S6.**
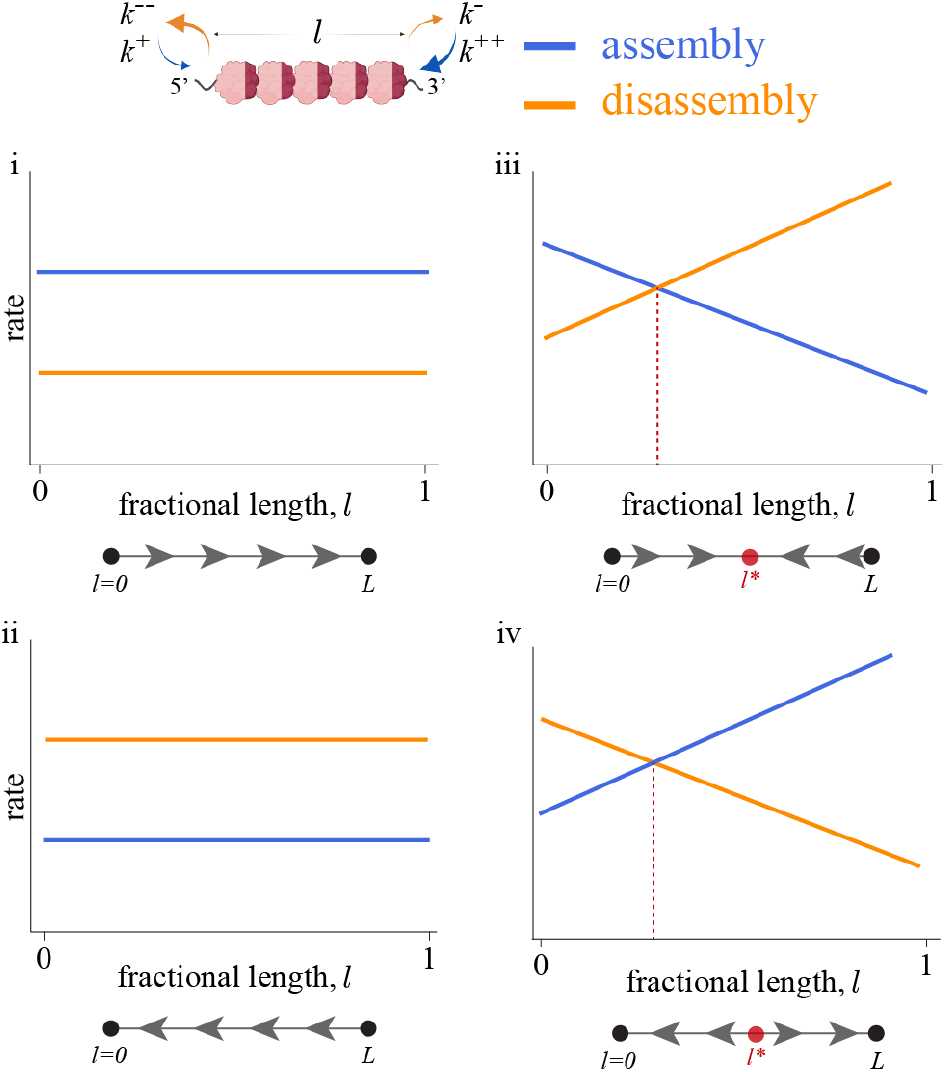
Homeostatic balance of RecA filament length arising from RecN-influenced assembly-disassembly of RecA units on the DNA strand. In order to obtain a fixed mean filament length by assembly-disassembly, the rates have to be dependent on the length *l.* The corresponding flow diagrams are shown below each panel.

Two final observations from the experiments: (i) The RecA filament appears to always nucleate from the DSB site, and does not seem to nucleate elsewhere on the DNA strand. This suggests that the nucleation is strongly heterogeneous. (ii) Since the filament integrity is maintained, i.e., the filament does not exhibit any breaks during its traversal across the cell, we assume that the RecA-RecN assembles/disassembles only at the two ends of the filament and not in the interior.

#### C. Stochastic Dynamics of the RecA-RecN filament

As discussed in the main text, our observation that the RecN-assisted movement of the RecA filament is strongly correlated with the stochastic remodelling of its length, immediately suggests that the RecN ATPase drives the RecA filament both by its motor activity via extrusion and the assembly-disassembly rates of RecA units on the DNA strand. Here we describe a stochastic kinetic model that integrates both the RecN-assisted DNA mobility and the dynamics of RecN influenced assembly-disassembly of the RecA units on the DNA strand using the effective reaction network described above.

We describe the stochastic dynamics of nucleation, growth and movement of the filament using a kinetic approach. If the movement arises from a combination of motor-assisted movement and assembly-disassembly processes, it is then appropriate to describe the RecA dynamics in the lab frame, taking into account both the movement of the DNA strand and the movement of the RecA filament on the DNA strand (Fig. S7).

Nucleation occurs at the DSB site, with a binding rate *k*^+^. As more RecA units are added, it grows into a RecA-filament with a left end x_*r*_ and right end x_*r*_. As a result of the stochastic assembly-disassembly processes, the ends x_*r*_ and x_*r*_ move to the left or right with specific rates.

We denote by *k*^-^ the bare rate of unbinding of the RecA-RecN complex. Initially, we expect *k*^+^ > *k*^-^, for the RecA-RecN to form a nascent filament post nucleation. Subsequently, since the filament integrity is maintained, we will assume that the RecA-RecN assembles/disassembles only at the two ends, x_*l*_ and x_*r*_, of the filament Fig. S7. To account for the RecN-induced unidirectional movement of the filament in the wild type cells, we allow for a different assembly rate at the right, *k*^++^, and a disassembly rate at the left, *k*^--^.

Mathematically, the pure motor-driven extrusion limit corresponds to a perfect correlation between *k*^++^ and *k*^--^ processes. For simplicity, we will use the language of assembly-disassembly processes, but it should be borne in mind that the stochastic dynamics encompasses *both* motor-assisted movement and assembly-disassembly and their combinations.

Since we are concerned with the dynamics of the filament well away from the cell poles, we have assumed that the cell boundaries do not affect the dynamics of x_*l*_ and x_*r*_.

**FIG. S7.**
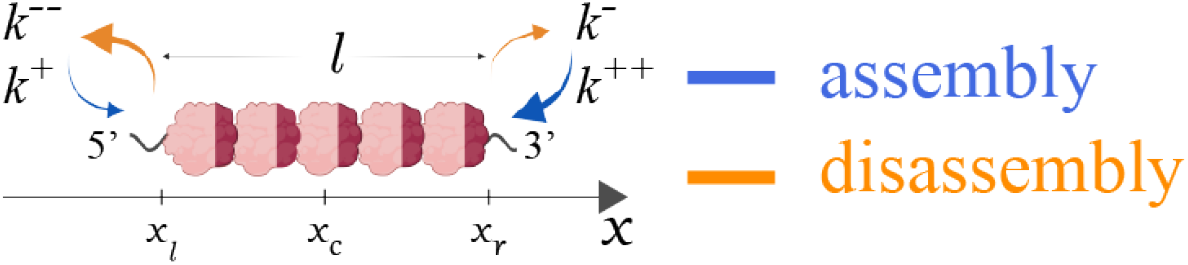
The RecN-assisted RecA filement dynamics relative to the lab frame (x-axis) is a combination of motor-assisted movement and assembly-disassembly.

Every time a RecA-RecN monomer is added or removed from the right end, x_*r*_ moves to the right or left by an amount *σ* (similarly for the left end x_*l*_). The right and left ends of the RecA-RecN filament thus move as a biased random walker with a stochastic dynamics given by the Langevin equation,

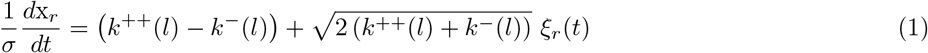

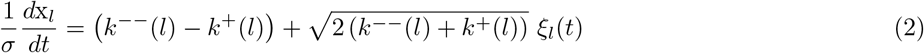

which comprises a drift term

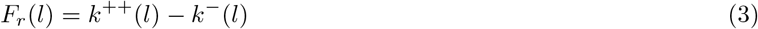

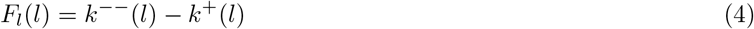

and a diffusive term,

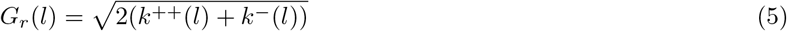

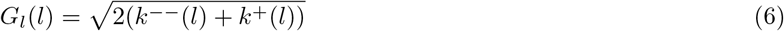

that multiplies the noises *ξ_r_*(*t*) and *ξ_l_*(*t*). The noise is taken to be a Gaussian white noise with zero mean and unit variance. We may rewrite Eq. 1,2 in terms of the the centroid and filament length, 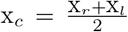 and *l* = x_*r*_ – x_*l*_, respectively,

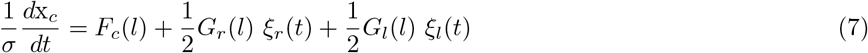

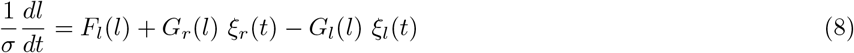

where

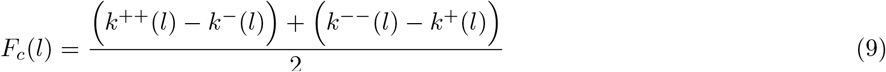

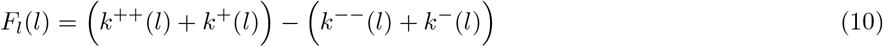

We will henceforth remember to measure x_*c*_ and *l* in units of *σ*, and so they will now be considered dimensionless. We note two features of the above equations -

1. Eq. 7 for the centroid x_*c*_ is a function of *l* and noise alone.
2. Eq. 8 for the length *l* is independent of x_*c*_.

Since the rates and hence *G_r_,G_l_* are dependent on the stochastic variable *l*, the noise is *multiplicative,* and hence special care needs to be taken to interpret the stochastic equation [15]. We will choose the Stratonovich interpretation.

These observations imply that one can obtain an expression for the mean centroid velocity 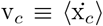, by simply averaging Eq. 7 over all noise realisations,

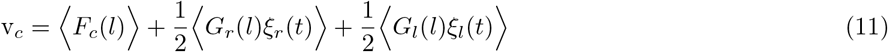

The last two averages on the right side of the above equation are nonzero due to the Stratonovich interpretation. To evaluate these, we make use of Novikov’s theorem for gaussian noise *ξ* with zero mean, which takes the form [18],

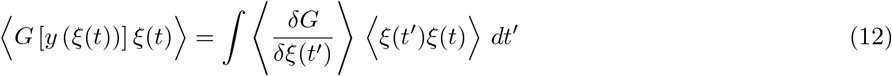

For a noise *ξ* with unit variance,

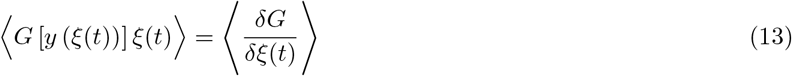

Using Novikov’s theorem, we obtain

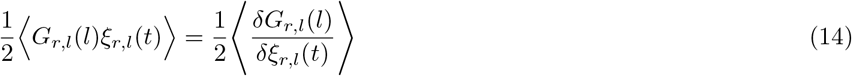

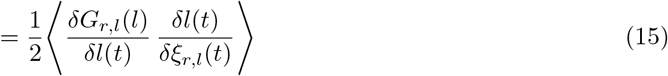

by the chain rule. The response function, *δl*(*t*)/*δξ_rl_*(*t*), is calculated from Eq. 8, using the Stratonovich interpretation,

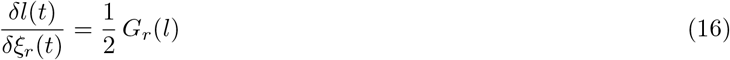

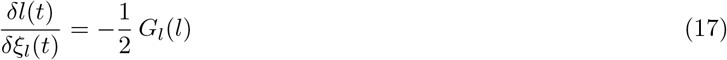

These results from Novikov’s theorem allow us to evaluate the averages in Eq. 11, to obtain,

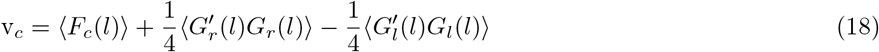

where prime denotes a derivative with respect to *l*. To evaluate these moments, for prescribed functions *F_c_*(*l*) and *G_r,l_*(*l*), we need to obtain the marginal distribution *P*(*l,t*). This can be obtained by constructing a Fokker-Planck equation for Eq. 8.

To do this, we first rewrite Eq. 8 as,

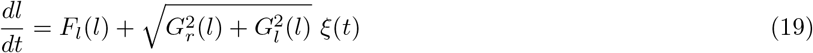

where *ξ*(*t*) = *ξ_r_*(*t*) + *ξ_l_*(*t*). The associated Fokker-Planck equation [15, 17] in the Stratonovich interpretation (ref) is,

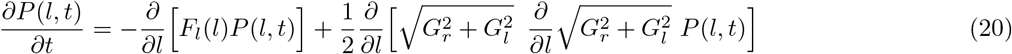

which is of the form,

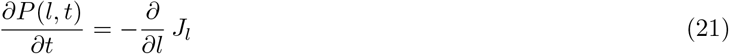

We will be interested in the stationary solution of Eq. 20 at steady state, i.e., when *∂P*(*l,t*)/*∂t* = 0. Further, since *l* is bounded, the probability current *J_l_* = 0, i.e.,

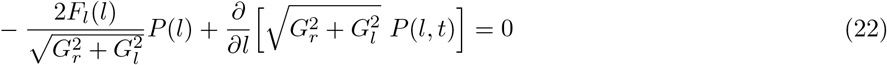

The solution obtained by straightforward integration is,

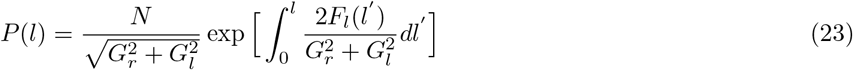

where *N* is obtained from the normalization condition,

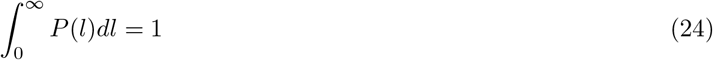

Eqs. 18 and 23 together with the normalization condition, are the main results of this section. To compare with the experimental observations in wild type and *ΔrecN* cells, we need to provide the functional forms of F¿ and *G_r,l_*.

But before doing that, we can make some general comments. From Eq. 10, we see that *F_l_* is the net imbalance between assembly and diassembly, and so *l* will grow if *F_l_* > 0, will shrink if *F_l_* < 0, and a balance is met when *F_l_* = 0. To obtain a distribution that is peaked at a finite *l*, one must have *F*(*l*) > 0 at small *l* (assembly dominates disassembly) and *F_l_* < 0 at large *l* (disassembly dominates assembly). The steady state distribution *P*(*l*) will then peak at *l*\*, given by the balance condition *F_l_* = 0. This distribution will typically be skewed about its peak value.

### S3. COMPARISON WITH EXPERIMENTS : EXTRACTING FITTING PARAMETERS

#### A. Wild type cells

Following the discussion in Sect.S2B, we take the RecN-dependent disassembly rate *k*^--^ at the left end of the filament to be linearly dependent on *l* and all other rates to be constant, i.e.,

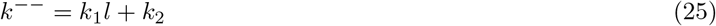

This implies that for large *l*, disassembly exceeds assembly and provides a mechanism for homeostatic balance at a finite fixed point *l*\*.

Such negative feedback control by RecN could be direct, via influence of RecN on RecA turnover within the filament or indirect via modulation of the underlying DNA substrate [6, 7, 22]. This could drive RecA turnover at opposite ends of the filament [3, 5, 20, 21].

Using this form in Eqs. 5, 6, 4 we get,

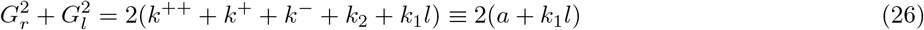

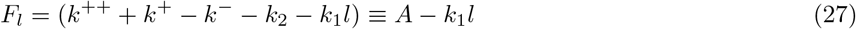

With these forms, the steady state distribution (Eq. 23) is,

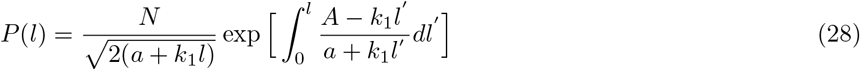

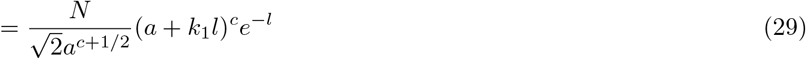

where,

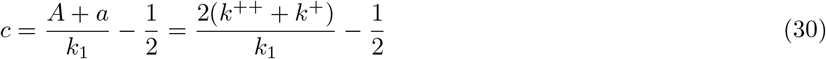

**FIG. S8.**
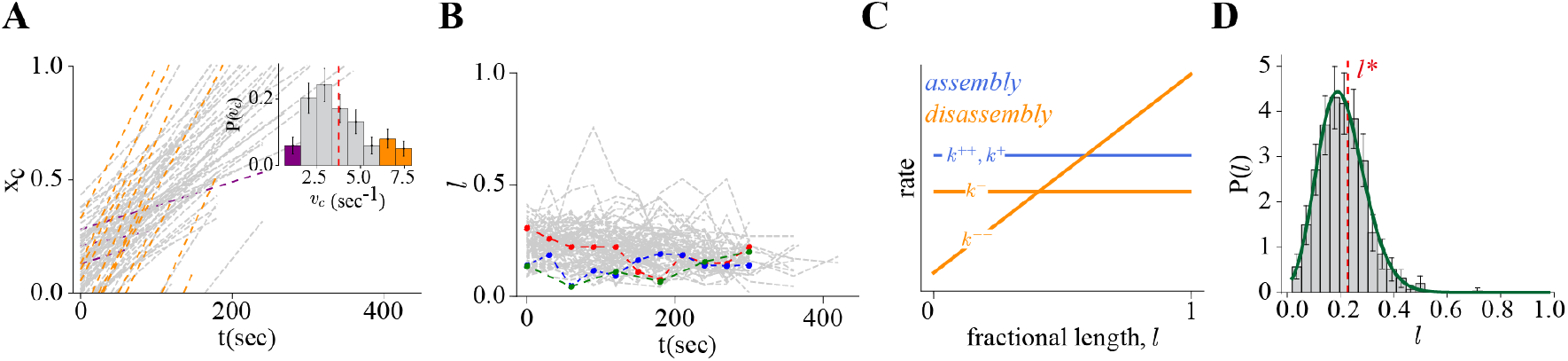
(A) Centroid position x_*c*_ versus time for each wild type cell at steady state. The centroid positions are fit to straight lines using a least-squares fit, from which we extract (inset) the distribution of the centroid velocity P (v_*c*_), which is sharply peaked at (v_*c*_) = 0.004 ± 0.002. The purple and orange data are outliers. (B) Fractional filament length *l* for each wild type cell versus time at steady state. The coloured dots and lines denote representative time series for *l*. (C) The disassembly rate (*k*) is taken to be length dependent while the assembly rates (*k*^++^, *k*^+^) are constant. (D) Fitting the experimental data (gray histogram) with the theory Eq. 32 (green). Red dashed vertical line is the mean length *l*\*.

At this stage we comment on some robust qualitative features of the derived distribution P(*l*), which are readily apparent in the observed wild type distribution. skewness, 〈*l*〈(mean *l* for a single traversal)> *l*\*, linearity for small *l*, exponential decay for large *l*,

We determine *N* from the normalization condition, Eq. 24, giving

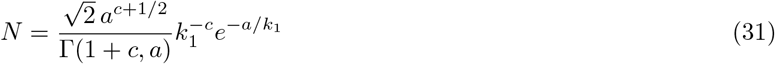

where the argument of the incomplete Γ function, *c* + 1 > 0, and the normalised distribution, (recall that *l* is in units of *σ*),

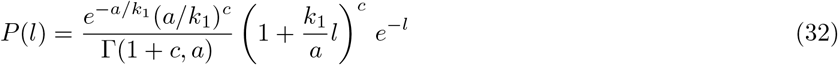

The qualitative features of the observed wild type distribution, namely (i) *P*(*l* → 0) > 0, (ii) *P*(*l*) is linear in *l* for small *l*, (iii) *P*(*l*) is a skewed distribution about the peak, (iv) The peak value of *l* is less than the mean, i.e., *l*\* < (l) and (v) *P*(*l*) falls off exponentially for larger *l*, are features of the distribution Eq. 32.

We find that the normalized form of (Eq. 32), fits the observed wild type distribution perfectly (Fig. S8), with fit parameters, the prefactor

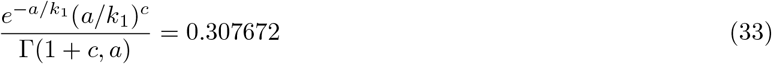

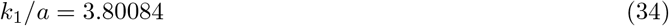

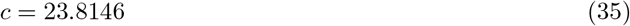

(Note that the experimental distribution in Fig. S8D is plotted with the fractional length relative to the cell size, *l/L*. Taking this into account introduces another fit parameter in the exponential term in Eq. 32, which we determine to be *d* = 54.1388.) That is 3 independent parameters, *k*_1_, *a, c*, which can be written in terms of the 5 original kinetic parameters *k*^+^, *k*^-^, *k*^++^, *k*_1_, *k*_2_, implying there is a lot of degeneracy in the values of these kinetic parameters. This would suggest that the distribution is robust to slight cell-to-cell variations in these kinetic parameters. We verify this robustness by pooling the WT data sets into batches of 25 cells, and comparing their respective distributions (Fig. S1E).

Having determined *P*(*l*), we can now unpack the relation Eq. 18 between the mean centroid velocity v_*c*_ and the moments of *P*(*l*). Using Eqs. 9, 5, 6, we see that

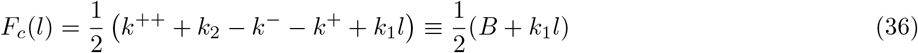

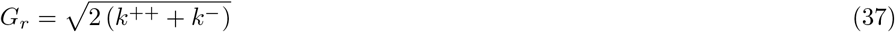

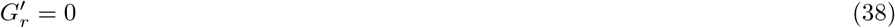

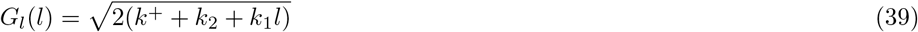

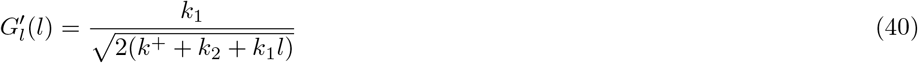

Thus,

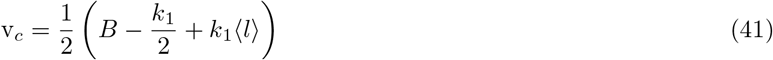

i.e., a linear relation between the mean centroid velocity and the mean filament length. This remarkable relation is verified in Fig. 3E [inset] in the main manuscript. We note that this graph in Fig. 3E, allows us to estimate a microscopic rate *k*_1_, the feedback associated with the length dependent disassembly rate. We also find that the intercept is negative, which gives a bound

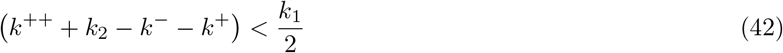

for the microscopic rates.

Following the discussion in Sect.S2B for the mutant Δ*recN* cells, we take the disassembly and assembly rates *k*^-^, *k*^++^ at the right end of the filament to be length *l* dependent and of the form,

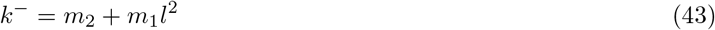

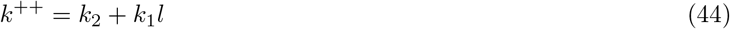

and all other rates to be constant. Here again for large *l*, disassembly exceeds assembly, leading to homeostatic balance at a finite fixed point *l*\*.

A possible molecular mechanism for such negative feedback control could be due to strain-dependent dissociation of RecA molecules at large filament lengths [23]. Indeed, evidence for cooperative/ allosteric loading of RecA has also been reported previously [4, 21, 23].

Using this form in Eqs. 5, 6, 4 we get,

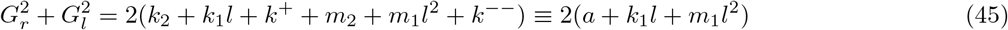

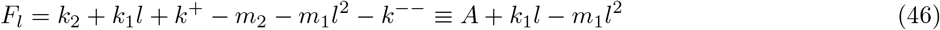

With these forms, the steady state distribution (Eq. 23) is,

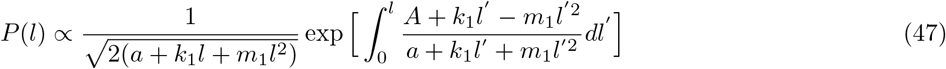

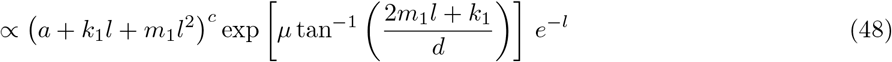

where, the parameters are known functions of the rates.

**FIG. S9.**
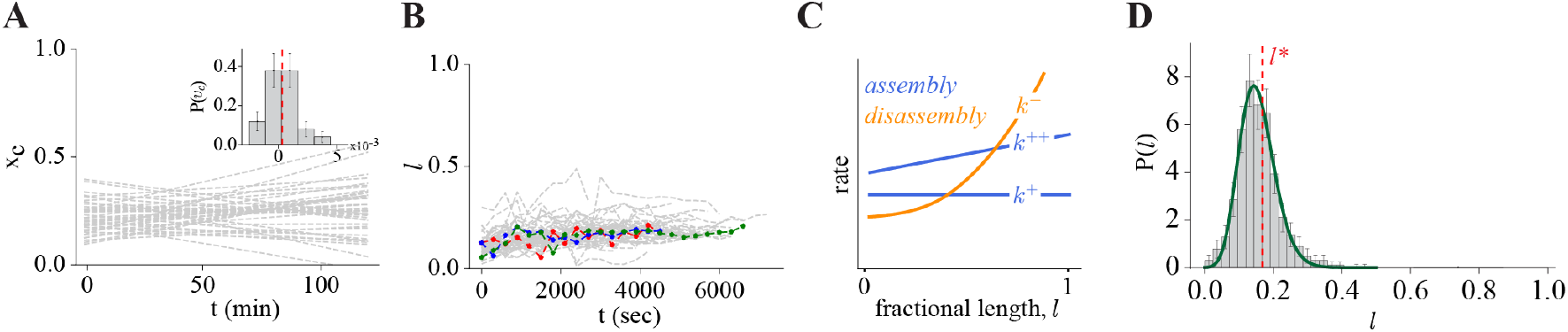
(A) Centroid position xc versus time for each wild type cell at steady state. The centroid positions are fit to straight lines using a least-squares fit, from which we extract (inset) the distribution of the centroid velocity P (v_*c*_), which is sharply peaked at (v_*c*_) = 0.0004 ± 0.00005. The purple and orange data are outliers. (B) Fractional filament length *l* for each wild type cell versus time at steady state. The coloured dots and lines denote representative time series for *l*. (C) The assembly rate (*k*^++^ = –*k*_1_*l*^2^ + *k*_2_) is taken to be length dependent while the disassembly rates (*k*^--^, *k*^-^) are constant. (D) Fitting the experimental data (gray histogram) with the theory curve (green). Red dashed vertical line denotes the mean length *l*\*.

We find that the distribution Eq. 48, when normalised numerically, fits the observed *ΔrecN* distribution perfectly (Fig. S9D), with normalisation constant, *A* = 5.3453, the rest of the parameters

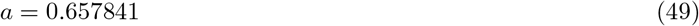

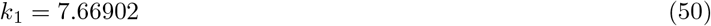

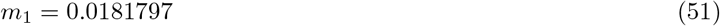

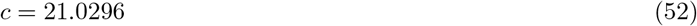

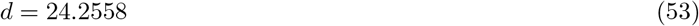

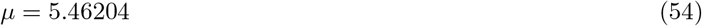

and the parameter that enters the exponential term *δ* = 91.9232 (since distribution is plotted in terms of the fractional length).

If we assume upfront that in *ΔrecN* cells, left end of the growing filament does not change from its initial position, then the stochastic dynamics of the filament only involves the Langevin equation for x_*r*_, Eq. 1. The filament length *l* = x_*r*_, while the centroid position 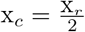 changes only because of the filament elongation.

---

**Supplementary table 1:** Strains used in present study.

| Strain name | Genotype | Strain construction/ source |
| --- | --- | --- |
| CB15N | NA1000 |  |
| NABC439 | CB15N; $\Delta recN::gent^R$ | (<sup>1</sup>) |
| NABC2 | CB15N; $\Delta recA$ | (<sup>2</sup>) |
| NABC443 | CB15N; $\Delta recA::gent^R$ | NABC2 was transformed with pNABC496 followed by pNABC482 to link *recA* deletion with *gent^R*. |
| NABC444 | CB15N; $\Delta recA::kan^R$ | NABC2 was transformed with pNABC496 followed by pNABC497 to link *recA* deletion with *kan^R*. |
| NABC445 | CB15N; $P_{xyl}-recA-mCherry::spec^R$ | CB15N was transformed with pNABC484 plasmid to integrate *recA-mCherry* linked to *spec^R* at the $P_{xyl}$ locus. |
| NABC446 | CB15N; $P_{xyl}-recA-mCherry::gent^R$ | CB15N was transformed with pNABC481 plasmid to integrate *recA-mCherry* linked to *gent^R* at the $P_{xyl}$ locus. |
| NABC447 | CB15N; $P_{xyl}-recA-YFP::gent^R$ | CB15N was transformed with pNABC487 plasmid to integrate *recA-YFP* linked to *gent^R* at the $P_{xyl}$ locus. |
| NABC448 | CB15N; $P_{xyl}-recA-YFP::spec^R$ | CB15N was transformed with pNABC480 plasmid to integrate *recA-YFP* linked to *spec^R* at the $P_{xyl}$ locus. |
| NABC449 | CB15N; $recN-YFP::spec^R$ | CB15N was transformed with pNABC489 plasmid to integrate *recN-YFP* linked to *spec^R* at the endogenous locus through two-step recombination procedure (<sup>3</sup>). |
| NABC450 | CB15N; $mipZ-mCherry::kan^R$ | CB15N was transformed with pNABC485 plasmid to integrate *mipZ-mCherry* linked to *kan^R* at the endogenous locus. |
| NABC451 | CB15N; $mipZ-mCherry::gent^R$ | CB15N was transformed with pNABC486 plasmid to integrate *mipZ-mCherry* linked to *gent^R* at the endogenous locus. |
| NABC452 | CB15N; $recN(D472A)::spec^R$ | CB15N was transformed with pNABC490 plasmid to generate a specific mutation in the ATP binding domain of RecN through two- step recombination procedure. The resultant strain was transformed with pNABC488 to link *recN(D472A)* with *spec^R*. |
| NABC453 | CB15N; $recN(E473Q)::spec^R$ | CB15N was transformed with pNABC491 plasmid to generate a specific mutation in the ATP hydrolysis domain of RecN through two- step recombination procedure. The resultant strain was transformed with pNABC488 to link *recN(E473Q)* with *spec^R*. |
| NABC454 | CB15N; $pMT687-P_{sidA}-YFP::kan^R$ | CB15N was transformed with pNABC498 plasmid. |
| NABC455 | CB15N; $P_{xyl}$-*recA*-*mCherry::gent<sup>R</sup>*; pMT687-$P_{sidA}$-*YFP::kan<sup>R</sup>* | NABC446 was transformed with pNABC498 plasmid. |
| NABC456 | CB15N; Δ*recA::gent<sup>R</sup>*; *recN*-*YFP::spec<sup>R</sup>* | NABC449 was transduced with lysate of a strain harboring *recA* deletion (NABC443) and selected for *gent<sup>R</sup>* that is linked to deletion. |
| NABC457 | CB15N; Δ*recN::gent<sup>R</sup>*; $P_{xyl}$-*recN*-*YFP::spec<sup>R</sup>* | NABC439 was transformed with pNABC483 plasmid to integrate *recN*-*YFP* linked to *spec<sup>R</sup>* at the $P_{xyl}$ locus. |
| NABC458 | CB15N; Δ*recN::gent<sup>R</sup>*; $P_{xyl}$-*recN*-*YFP::spec<sup>R</sup>*; Δ*recA::kan<sup>R</sup>* | NABC457 was transduced with lysate of a strain harboring *recA* deletion (NABC444) and selected for *kan<sup>R</sup>* that is linked to deletion. |
| NABC459 | CB15N; Δ*recN::gent<sup>R</sup>*; pMT687-$P_{sidA}$-*YFP::kan<sup>R</sup>* | NABC439 was transformed with pNABC498 plasmid. |
| NABC460 | CB15N; $P_{xyl}$-*recA*-*mCherry::spec<sup>R</sup>*; Δ*recN::gent<sup>R</sup>* | NABC445 was transduced with lysate of a strain harboring *recN* deletion (NABC439) and selected for *gent<sup>R</sup>* that is linked to deletion. |
| NABC413 | $P_{lacI}$-*lacI*(*hfa* locus); $P_{lac}$-*dnaA*(*dnaA* locus) | (<sup>1</sup>) |
| NABC461 | $P_{lacI}$-*lacI*(*hfa* locus); $P_{lac}$-*dnaA*(*dnaA* locus); I-SceI site(after CCNA_00029)::*tet<sup>R</sup>*; $P_{van}$-*I-SceI::chlor<sup>R</sup>* | (<sup>4</sup>) |
| NABC462 | $P_{lacI}$-*lacI*(*hfa* locus); $P_{lac}$-*dnaA*(*dnaA* locus); I-SceI site(after CCNA_00029)::*tet<sup>R</sup>*; $P_{van}$-*I-SceI::chlor<sup>R</sup>*; Δ*recA::gent<sup>R</sup>* | NABC461 was transduced with lysate of a strain harboring *recA* deletion (NABC443) and selected for *gent<sup>R</sup>* that is linked to deletion. |
| NABC463 | $P_{lacI}$-*lacI*(*hfa* locus); $P_{lac}$-*dnaA*(*dnaA* locus); I-SceI site(after CCNA_00029)::*tet<sup>R</sup>*; $P_{van}$-*I-SceI::chlor<sup>R</sup>*; Δ*recN::gent<sup>R</sup>* | NABC461 was transduced with lysate of a strain harboring *recN* deletion (NABC439) and selected for *gent<sup>R</sup>* that is linked to deletion. |
| NABC464 | $P_{lacI}$-*lacI*(*hfa* locus); $P_{lac}$-*dnaA*(*dnaA* locus); I-SceI site(after CCNA_00029)::*tet<sup>R</sup>*; $P_{van}$-*I-SceI::chlor<sup>R</sup>*; Δ*smc::kan<sup>R</sup>* | NABC461 was transduced with lysate of a strain harboring *smc* deletion (Tung B. K. Le, 2013) and selected for *kan<sup>R</sup>* that is linked to deletion. |
| NABC465 | $P_{lacI}$-*lacI*(*hfa* locus); $P_{lac}$-*dnaA*(*dnaA* locus); I-SceI site(after CCNA_00029)::*tet<sup>R</sup>*; $P_{van}$-*I-SceI::chlor<sup>R</sup>*; *mipZ*-*YFP::kan<sup>R</sup>* | (<sup>4</sup>) |
| NABC466 | $P_{lacI}$-*lacI*(*hfa* locus); $P_{lac}$-*dnaA*(*dnaA* locus); I-SceI site(after CCNA_00029)::*tet<sup>R</sup>*; $P_{van}$-*I-SceI::chlor<sup>R</sup>*; *mipZ*-*YFP::kan<sup>R</sup>*; Δ*recA::gent<sup>R</sup>* | (<sup>4</sup>) |
| NABC467 | $P_{lacI}$-*lacI*(*hfa* locus); $P_{lac}$-*dnaA*(*dnaA* locus); I-SceI site(after CCNA_00029)::*tet<sup>R</sup>*; $P_{van}$-*I-SceI::chlor<sup>R</sup>*; *mipZ*-*mCherry::kan<sup>R</sup>*; $P_{xyl}$-*recA*-*YFP::gent<sup>R</sup>* | NABC461 was transduced with lysate of a strain harboring *mipZ*-*mCherry* linked to *kan<sup>R</sup>* (NABC450) integrated at endogenous locus. Resultant strain was transduced with lysate of a strain harboring *recA*-*YFP* linked to *gent<sup>R</sup>* (NABC447) integrated at the $P_{xyl}$ locus. |
| NABC468 | *P<sub>lacI</sub>-lacI(hfa locus)*; *P<sub>lac</sub>-dnaA(dnaA locus)*; I-SceI site(after CCNA_00029)::*tet<sup>R</sup>*; *P<sub>van</sub>-I-SceI::chlor<sup>R</sup>*; *mipZ-mCherry::kan<sup>R</sup>*; *P<sub>xyI</sub>-recA-YFP::spec<sup>R</sup>*; *ΔrecN::gent<sup>R</sup>* | NABC461 was transduced with lysate of a strain harboring *mipZ-mCherry* linked to *kan<sup>R</sup>* (NABC450) integrated at endogenous locus. Resultant strain was transduced with lysate of a strain harboring *recA-YFP* linked to *spec<sup>R</sup>* (NABC448) integrated at the *P<sub>xyI</sub>* locus. Finally, lysate of NABC439 was transduced to generate a *recN* deletion linked to *gent<sup>R</sup>*. |
| NABC469 | *P<sub>lacI</sub>-lacI(hfa locus)*; *P<sub>lac</sub>-dnaA(dnaA locus)*; I-SceI site(after CCNA_00029)::*tet<sup>R</sup>*; *P<sub>van</sub>-I-SceI::chlor<sup>R</sup>*; *mipZ-mCherry::kan<sup>R</sup>*; *P<sub>xyI</sub>-recA-YFP::gent<sup>R</sup>*; *recN(D472A)::spec<sup>R</sup>* | NABC467 was transduced with lysate of a strain harboring *recN(D472A)* linked to *spec<sup>R</sup>* (NABC452). |
| NABC470 | *P<sub>lacI</sub>-lacI(hfa locus)*; *P<sub>lac</sub>-dnaA(dnaA locus)*; I-SceI site(after CCNA_00029)::*tet<sup>R</sup>*; *P<sub>van</sub>-I-SceI::chlor<sup>R</sup>*; *mipZ-mCherry::kan<sup>R</sup>*; *P<sub>xyI</sub>-recA-YFP::gent<sup>R</sup>*; *recN(E473Q)::spec<sup>R</sup>* | NABC467 was transduced with lysate of a strain harboring *recN(E473Q)* linked to *spec<sup>R</sup>* (NABC453). |
| NABC471 | *P<sub>lacI</sub>-lacI(hfa locus)*; *P<sub>lac</sub>-dnaA(dnaA locus)*; I-SceI site(after CCNA_00727)::*tet<sup>R</sup>* | NABC413 was transformed with a pNPTS138 vector (<sup>5</sup>) to integrate *I-SceI* site after CCNA_00727 through two-step recombination procedure. |
| NABC472 | *P<sub>lacI</sub>-lacI(hfa locus)*; *P<sub>lac</sub>-dnaA(dnaA locus)*; I-SceI site(after CCNA_00727)::*tet<sup>R</sup>*; parSpMT1 site (after CCNA_00747)::*spec<sup>R</sup>*; *P<sub>van</sub>-I-SceI::chlor<sup>R</sup>* | (<sup>5</sup>) |
| NABC473 | *P<sub>lacI</sub>-lacI(hfa locus)*; *P<sub>lac</sub>-dnaA(dnaA locus)*; I-SceI site(after CCNA_00727)::*tet<sup>R</sup>*; *P<sub>van</sub>-I-SceI::chlor<sup>R</sup>*; parSpMT1 site (after CCNA_00747)::*spec<sup>R</sup>*; *ΔrecA::gent<sup>R</sup>* | (<sup>5</sup>) |
| NABC474 | *P<sub>lacI</sub>-lacI(hfa locus)*; *P<sub>lac</sub>-dnaA(dnaA locus)*; I-SceI site(after CCNA_00727)::*tet<sup>R</sup>*; *P<sub>van</sub>-I-SceI::chlor<sup>R</sup>*; parSpMT1 site (after CCNA_00747)::*spec<sup>R</sup>*; *ΔrecN::gent<sup>R</sup>* | NABC472 was transduced with lysate of a strain harboring *recN* deletion (NABC439) and selected for *gent<sup>R</sup>* that is linked to deletion. |
| NABC475 | *P<sub>lacI</sub>-lacI(hfa locus)*; *P<sub>lac</sub>-dnaA(dnaA locus)*; I-SceI site(after CCNA_00727)::*tet<sup>R</sup>*; *P<sub>van</sub>-I-SceI::chlor<sup>R</sup>*; *P<sub>xyI</sub>-recA-YFP::spec<sup>R</sup>* | NABC471 was transduced with lysate of a strain harboring I-SceI enzyme (NABC472) linked to *chlor<sup>R</sup>* integrated at the *P<sub>van</sub>* locus. The resultant strain was transduced with lysate of a strain harboring *recA-YFP* (NABC448) linked to *spec<sup>R</sup>* integrated at the *P<sub>xyI</sub>* locus. |
| NABC476 | BTH101; pKT25; pUT18C | (<sup>5</sup>) |
| NABC477 | BTH101; pKT25-*addB::kan<sup>R</sup>*;<br>pUT18C-*addA::carb<sup>R</sup>* | (<sup>6</sup>) |
| NABC478 | BTH101; pKT25-*recA::kan<sup>R</sup>*;<br>pUT18C-*recN::carb<sup>R</sup>* | BTH101 was co-transformed with pNABC494 (<sup>6</sup>) and pNABC493 for the bacterial-two-hybrid assay. |
| NABC479 | BTH101; pKT25-*recN::kan<sup>R</sup>*;<br>pUT18C-*recA::carb<sup>R</sup>* | BTH101 was co-transformed with pNABC495 (<sup>6</sup>) and pNABC492 for the bacterial-two-hybrid assay. |
| NABC508 | CB15N; *P<sub>xyl</sub>-ssb-YFP::kan<sup>R</sup>* | CB15N was transformed with pNABC512 plasmid to integrate *ssb-YFP* linked to *kan<sup>R</sup>* at the *P<sub>xyl</sub>* locus. |
| NABC509 | *P<sub>lacI</sub>-lacI(hfa locus)*; *P<sub>lac</sub>-dnaA(dnaA locus)*; I-SceI site(after CCNA_00029)::*tet<sup>R</sup>*; *P<sub>van</sub>-I-SceI::chlor<sup>R</sup>*; *mipZ-mCherry::gent<sup>R</sup>*; *P<sub>xyl</sub>-ssb-YFP::kan<sup>R</sup>* | NABC461 was transduced with lysate of a strain harboring *ssb-YFP* linked to *kan<sup>R</sup>* (NABC508) integrated at *P<sub>xyl</sub>* locus. Resultant strain was transduced with lysate of a strain harboring *mipZ-mCherry* linked to *gent<sup>R</sup>* (NABC451) integrated at endogenous locus. Finally, lysate of NABC439 was transduced to generate a *recN* deletion linked to *gent<sup>R</sup>*. |
| NABC510 | CB15N; *mipZ-mCherry::spec<sup>R</sup>* | CB15N was transformed with pNABC513 plasmid to integrate *mipZ-mCherry* linked to *spec<sup>R</sup>* at the endogenous locus. |
| NABC511 | *P<sub>lacI</sub>-lacI(hfa locus)*; *P<sub>lac</sub>-dnaA(dnaA locus)*; I-SceI site(after CCNA_00029)::*tet<sup>R</sup>*; *P<sub>van</sub>-I-SceI::chlor<sup>R</sup>*; *mipZ-mCherry::spec<sup>R</sup>*; *P<sub>xyl</sub>-ssb-YFP::kan<sup>R</sup>*; *ΔrecN::gent<sup>R</sup>* | NABC461 was transduced with lysate of a strain harboring *ssb-YFP* linked to *kan<sup>R</sup>* (NABC508) integrated at *P<sub>xyl</sub>* locus. Resultant strain was transduced with lysate of a strain harboring *mipZ-mCherry* linked to *spec<sup>R</sup>* (NABC510) integrated at endogenous locus. Finally, lysate of NABC439 was transduced to generate a *recN* deletion linked to *gent<sup>R</sup>*. |
| NABC516 | *P<sub>lacI</sub>-lacI(hfa locus)*; *P<sub>lac</sub>-dnaA(dnaA locus)*; I-SceI site(after CCNA_00029)::*tet<sup>R</sup>*; *P<sub>van</sub>-I-SceI::chlor<sup>R</sup>*; *Δsmc::kan<sup>R</sup>*; *P<sub>xyl</sub>-recA-YFP::spec<sup>R</sup>* | NABC464 was transduced with a lysate harbouring *recA-YFP* (NABC448) linked to *spec<sup>R</sup>* integrated at the *P<sub>xyl</sub>* locus. |
| NABC517 | *P<sub>lacI</sub>-lacI(hfa locus)*; *P<sub>lac</sub>-dnaA(dnaA locus)*; I-SceI site(after CCNA_00029)::*tet<sup>R</sup>*; *P<sub>van</sub>-I-SceI::chlor<sup>R</sup>*; *P<sub>xyl</sub>-recA-mCherry::gent<sup>R</sup>*; *recN-YFP::spec<sup>R</sup>* | NABC461 was transduced with a lysate harbouring *recA-mCherry* (NABC446) linked to *gent<sup>R</sup>* integrated at the *P<sub>xyl</sub>* locus. Resultant strain was transduced with a lysate harbouring *recN-YFP* (NABC449) linked to *spec<sup>R</sup>* at the endogenous locus. |
| NABC518 | BTH101; pKT25-*recN(E473Q)::kan<sup>R</sup>*;<br>pUT18C-*recA::carb<sup>R</sup>* | BTH101 was co-transformed with pNABC495 (<sup>6</sup>) and pNABC521 for the bacterial-two-hybrid assay. |
| NABC519 | BTH101; pKT25-*recN(D472A)::kan<sup>R</sup>*;<br>pUT18C-*recA::carb<sup>R</sup>* | BTH101 was co-transformed with pNABC495 (<sup>6</sup>) and pNABC522 for the bacterial-two-hybrid assay. |
| NABC520 | CB15N; *ΔlexA::tet<sup>R</sup>*;<br>*ΔsidA::markerless* | (<sup>7</sup>) |
| NABC265 | CB15N; *ΔlexA::tet<sup>R</sup>*;<br>*ΔsidA::markerless*; *ΔrecA::kan<sup>R</sup>* | NABC520 was transduced with lysate of a strain harboring *recA* deletion (NABC444) and selected for *kan<sup>R</sup>* that is linked to deletion. |
| NABC586 | CB15N; *recN(E473Q)* | CB15N was transformed with pNABC491 plasmid to generate a specific mutation in the ATP hydrolysis domain of RecN through two-step recombination procedure. |
| NABC587 | CB15N; *recN(K35A/E473Q)::kan<sup>R</sup>* | NABC587 was transformed with pNABC584 plasmid to generate a specific mutation in the ATP hydrolysis domain of RecN through two-step recombination procedure. The resultant strain was transformed with pNABC585 to link *recN(K35A/E473Q)* with *kan<sup>R</sup>*. |
| NABC588 | *P<sub>lacI</sub>-lacI(hfa locus); P<sub>lac</sub>-dnaA(dnaA locus); I-SceI site(after CCNA_00029)::tet<sup>R</sup>; P<sub>van</sub>-I-SceI::chlor<sup>R</sup>; P<sub>xyl</sub>-recA-YFP::gent<sup>R</sup>;* | NABC461 was transduced with lysate of a strain harboring *P<sub>xyl</sub>-recA-YFP* (NABC447) and selected for *gent<sup>R</sup>*. |
| NABC589 | *P<sub>lacI</sub>-lacI(hfa locus); P<sub>lac</sub>-dnaA(dnaA locus); I-SceI site(after CCNA_00029)::tet<sup>R</sup>; P<sub>van</sub>-I-SceI::chlor<sup>R</sup>; P<sub>xyl</sub>-recA-YFP::gent<sup>R</sup>; *recN(K35A/E473Q)::kan<sup>R</sup>** | NABC588 was transduced with lysate of a strain harboring *recN(K35A/E473Q)* (NABC587) and selected for *kan<sup>R</sup>*. |

**Supplementary table 2:**
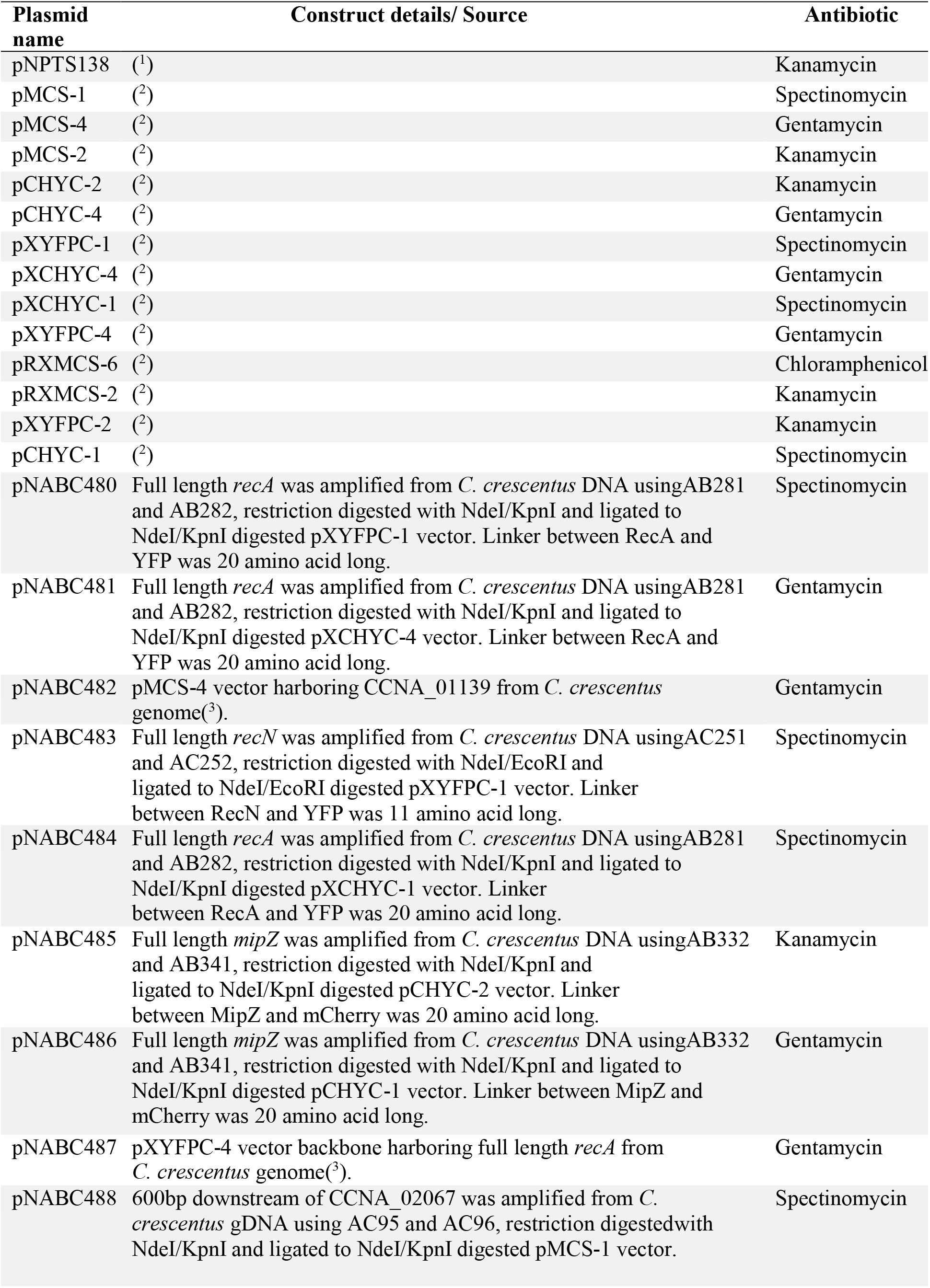
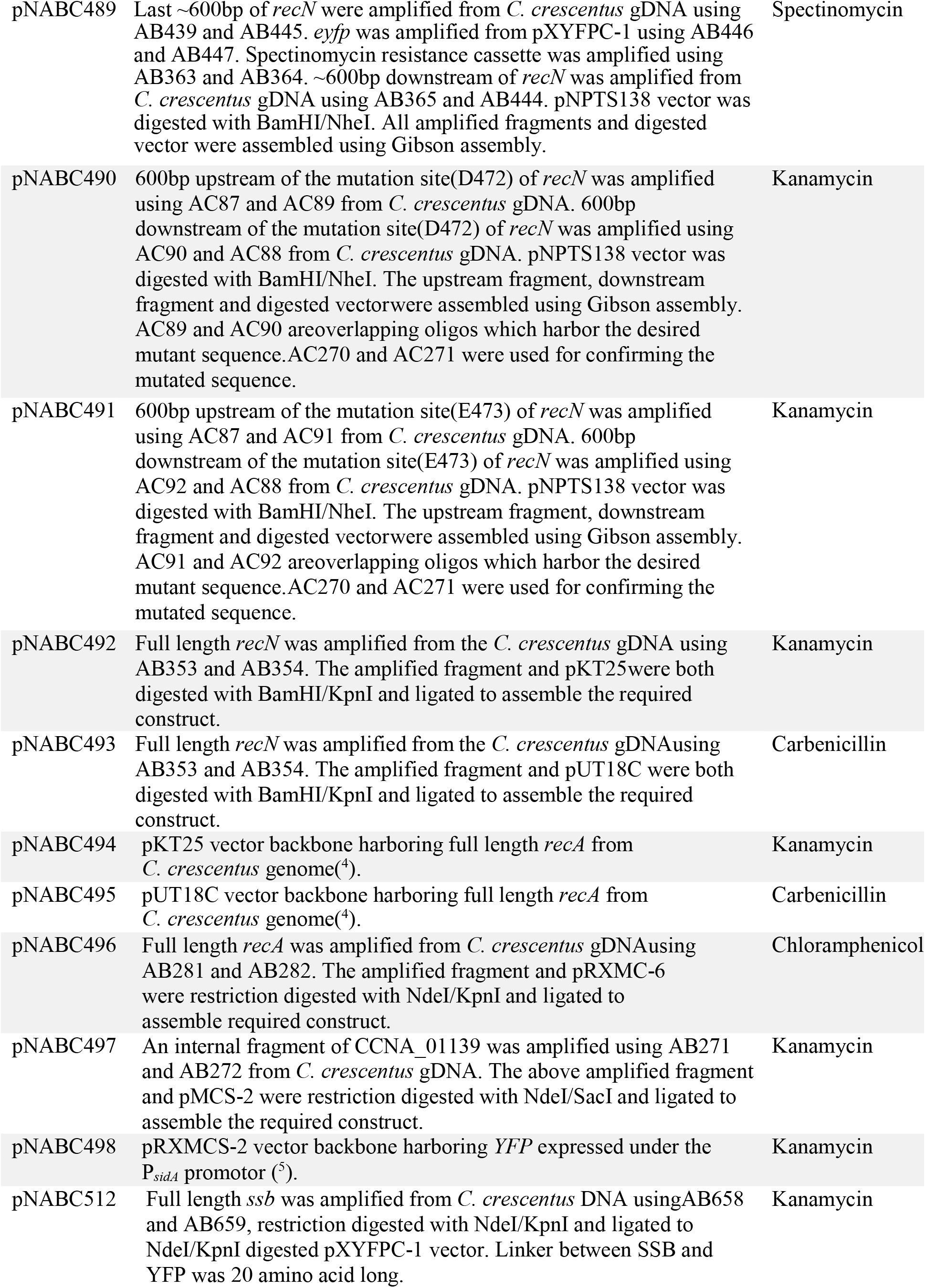
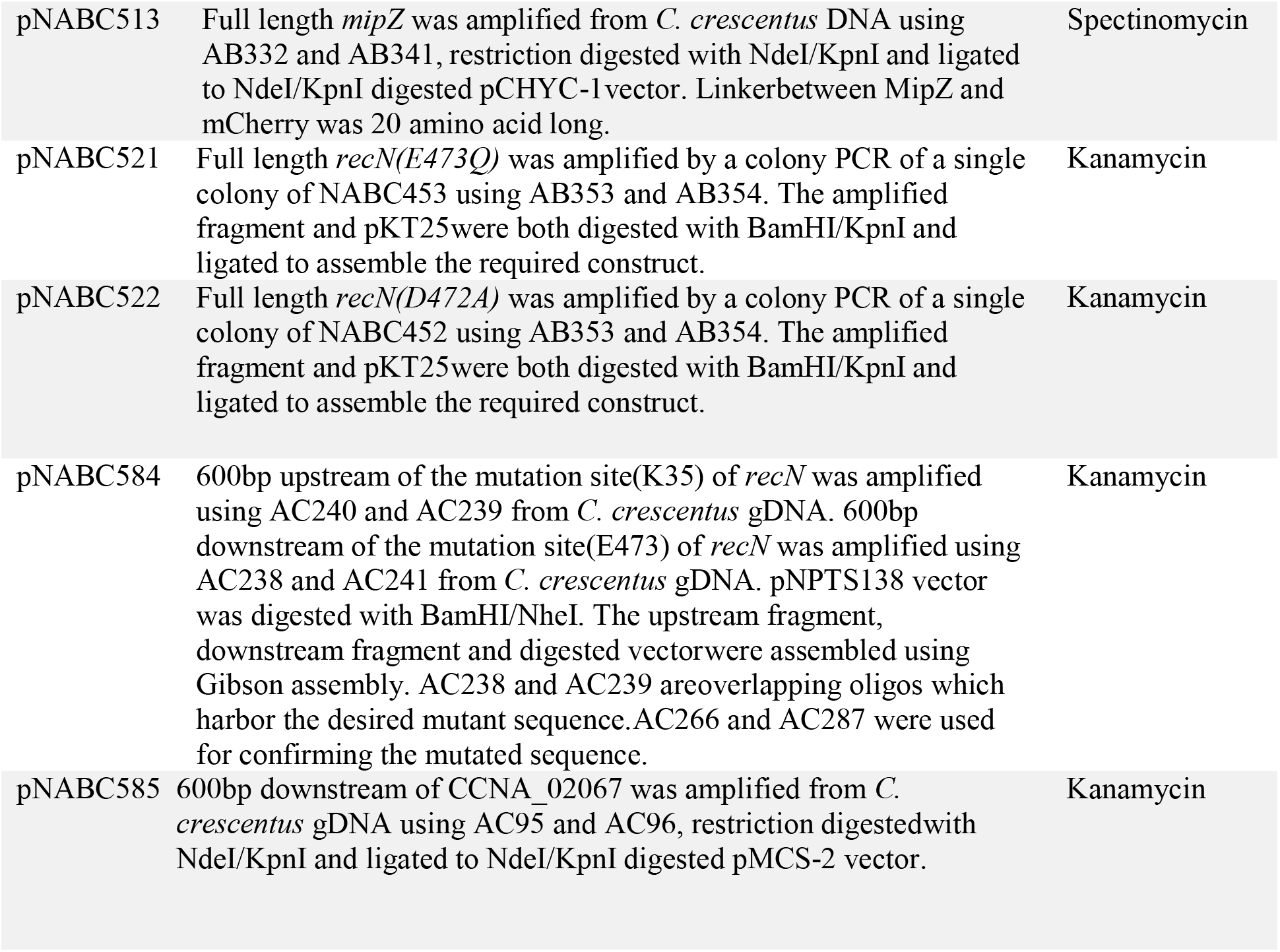
Plasmids used in present study.

**Supplementary table 3:**
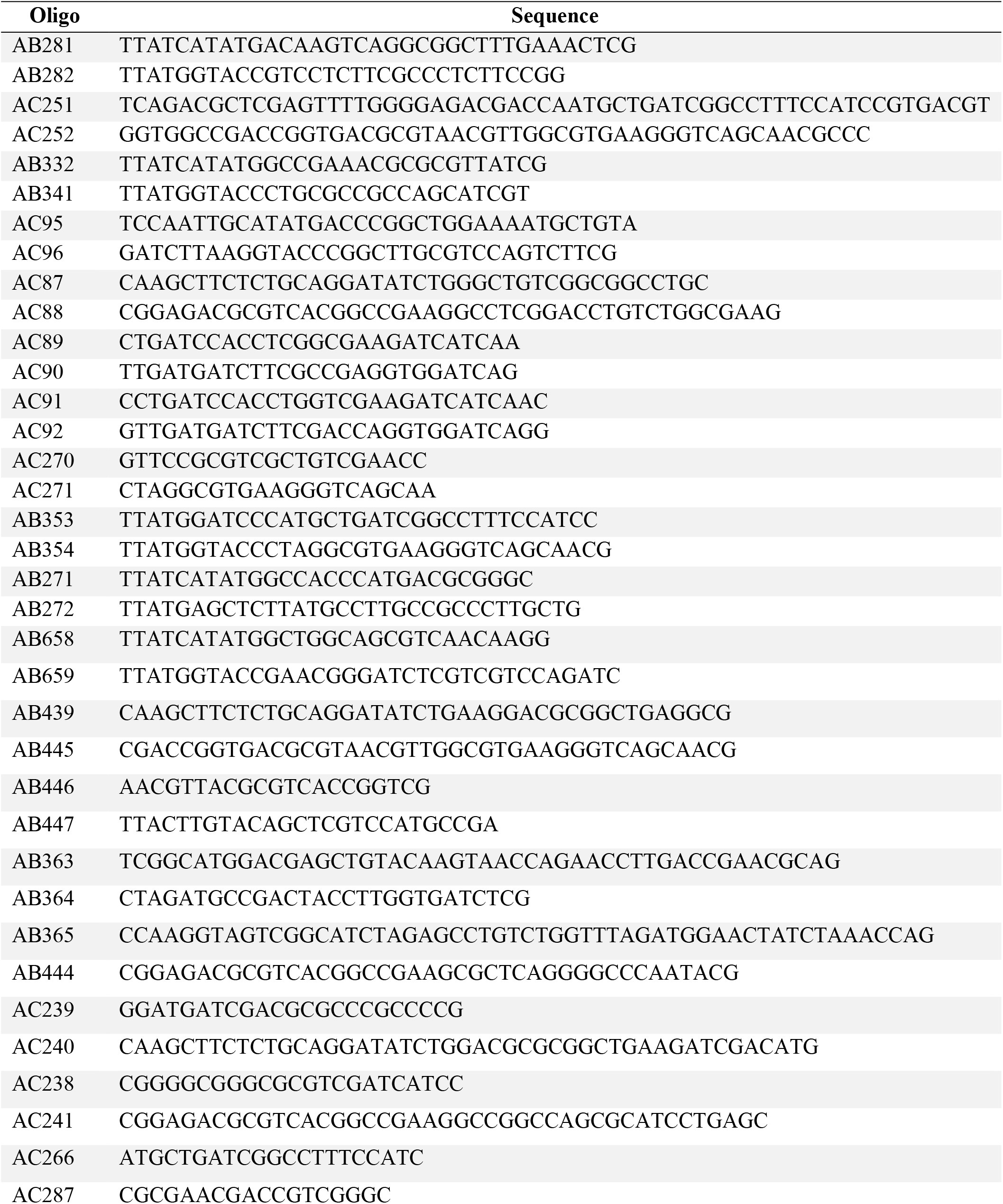
Oligos used in present study.

